# Longitudinal changes in brain metabolites in healthy subjects and patients with first episode psychosis (FEP): a 7-Tesla MRS study

**DOI:** 10.1101/2020.08.25.267419

**Authors:** Min Wang, Peter B. Barker, Nicola Cascella, Jennifer M. Coughlin, Gerald Nestadt, Frederick C. Nucifora, Thomas W. Sedlak, Alexandra Kelly, Laurent Younes, Donald Geman, Akira Sawa, Kun Yang

## Abstract

**Objective:** 7 Tesla (T) longitudinal magnetic resonance spectroscopy (MRS) offers a precise measurment of metabolic levels in human brain via a non-invasive approach. Studying longitudinal changes in neurometabolites could help identify trait and state markers for diseases and understand inconsistent findings from different researchers due to differences in the age of study participants and duration of illness. This study is the first to report novel longitudinal patterns in young adulthood from both physiological and pathological viewpoints using 7T MRS.

**Methods:** Utilizing a four-year longitudinal cohort with 38 first episode psychosis (FEP) patients (onset within 2 years) and 48 healthy controls (HC), the authors examined the annual percentage changes of 9 neurometabolites in 5 brain regions.

**Results:** Both FEP patients and HC subjects were found to have significant longitudinal reductions in glutamate (Glu) in the anterior cingulate cortex (ACC). Only FEP patients were found to have a significant decrease over time in γ-aminobutyric acid (GABA), N-acetyl aspartate (NAA), myo-inositol (mI), and total choline (tCho: phosphocholine plus glycerophosphocholine) in the ACC. Uniquely, glutathione (GSH) was found to have a near zero annual percentage change in both FEP patients and HC subjects in all 5 brain regions over a four-year timespan in young adulthood.

**Conclusions:** GSH could be a trait marker for diagnostic applications at least in young adulthood. Glu, GABA, NAA, mI, and tCho in the ACC are associated with the patient’s status and could be state markers for mechanistic studies of psychotic disorders, including those for progressive pathological changes and medication effects in young adulthood.

## Introduction

Proton magnetic resonance spectroscopy (MRS) allows for the non-invasive quantification of multiple compounds that are related to metabolism, neurotransmission and other processes in the human brain (1). It is a powerful technique, well appreciated in psychiatric studies to measure neurometabolic changes *in vivo* (2, 3). Compared to scans performed at lower field strengths, 7 Tesla (T) MRS offers a more precise measurement of metabolite levels (4–6). To date, there have only been a handful of 7T MRS studies using either first episode psychosis (FEP) (7) or schizophrenia (SZ) patients (8–13). However, all of these studies were cross-sectional and most of them used a fairly small number of subjects.

Inconsistent findings in neurometabolic changes have been observed in psychotic studies (14, 15). For example, some studies have found increased γ-aminobutyric acid (GABA) in the medial frontal cortex in SZ while other studies observed reductions (16). Several factors may cause these inconsistent findings such as the heterogeneity of the disease and possible differences in MRS techniques. One important factor could be the age of the study participants and their duration of illness. There is evidence that brain metabolites measured by 3T MRS change over time, either due to disease progression or in response to, or as a consequence of, therapeutic interventions (16–19). However, so far there is not a systematic study of the physiological and pathological effects on neurometabolic changes, especially during young adulthood (and late adolescence) when most patients experience FEP. The fact that only a few longitudinal MRS studies of FEP have been published (15, 17, 19–23), especially none to date performed at 7T, limits current progress in studying neurometabolic changes in early stages of psychotic disorders.

Another barrier for longitudinal studies is the statistical method. Most statistical methods require a relatively large sample size and multiple time points to obtain meaningful results (24). For example, the newly developed changepoint method could utilize longitudinal data to identify a significant time point that is related to the onset of disease. This approach has been successfully applied in the study of Alzheimer’s disease (25). However, this method cannot be used for studies where there are only 2 or 3 time points available. Commonly used approaches such as multivariate analysis of variance or regression models are also limited when only a small number of time points are available and missing data exists (26, 27). On the other hand, for practical reasons (e.g., time, budget, attrition), it is hard to acquire longitudinal cohorts with large sample sizes and multiple time points. A straightforward and easy analysis pipeline that can be used in studies with limited time points could reduce the barrier for longitudinal studies and boost new discoveries.

To fill these intellectual gaps, we now report a longitudinal study of FEP patients and healthy controls (HC) from physiological and pathophysiological viewpoints. This longitudinal study expands our previously published cross-sectional study of 7T MRS with 81 FEP patients and 91 HC subjects that reported a number of metabolic abnormalities in FEP patients (12): these included lower levels of GABA, glutamate (Glu), glutathione (GSH), and N-acetyl aspartate (NAA) in the anterior cingulate cortex (ACC) in FEP patients compared with HC subjects, decreased NAA in the thalamus and orbitofrontal region, and a reduction in GSH in the thalamus (12). Among the 172 participants in our published cross-sectional 7T MRS study (12), 38 FEP patients and 48 HC subjects returned for at least one follow-up evaluation during 4 years and represent the cohort in this longitudinal study. Through a new analytic pipeline, we observed longitudinal changes of key metabolites at both physiological and pathological levels in the present study.

## Methods

### Participants

This study was approved by the Johns Hopkins University School of Medicine Institutional Review Board and all subjects provided written informed consent. As stated in our previous publication (12), patients were within 24 months of the onset of positive symptoms at their baseline visit, as assessed by psychiatrists using the Structured Clinical Interview for DSM-IV (SCID) and medical records. Among the 172 participants in the previous cross-sectional study (12), 38 FEP patients and 48 HC subjects came back for yearly study visits, including 7T MRS scans, for up to 4 years.

### MR protocol

All participants were scanned using a 7T scanner (Philips ‘Achieva’, Best, Netherlands) equipped with a 32-channel receive head coil using a protocol previously described in detail (12). High-resolution (0.8 mm isotropic) T_1_-weighted anatomical images were acquired using an MPRAGE sequence. Spectra were recorded using the STEAM sequence (TR/TE/TM = 3000/14/33 ms, 128 excitations) with VAPOR water suppression from the thalamus (Thal, 20×30×15 mm^3^), orbitofrontal region (OFR; 20×20×20 mm^3^), anterior cingulate cortex (ACC; 30×20×20 mm^3^), dorsolateral prefrontal cortex (DLPFC; 25×20×20 mm^3^) and centrum semiovale (CSO; 40×20×15 mm^3^) (**Figure 1A**). A non-water suppressed acquisition was also collected with 2 excitations. **Figure 1B** shows representative spectra from the ACC of one subject at two time points, and the results of the spectral fitting routine. Prior to acquisition, field homogeneity was optimized up to the 2^nd^ order using the FASTMAP technique (28), and RF pulses calibrated using a localized power optimization scheme (29). Scan time was 6min 30s per voxel location.

**Figure 1.**
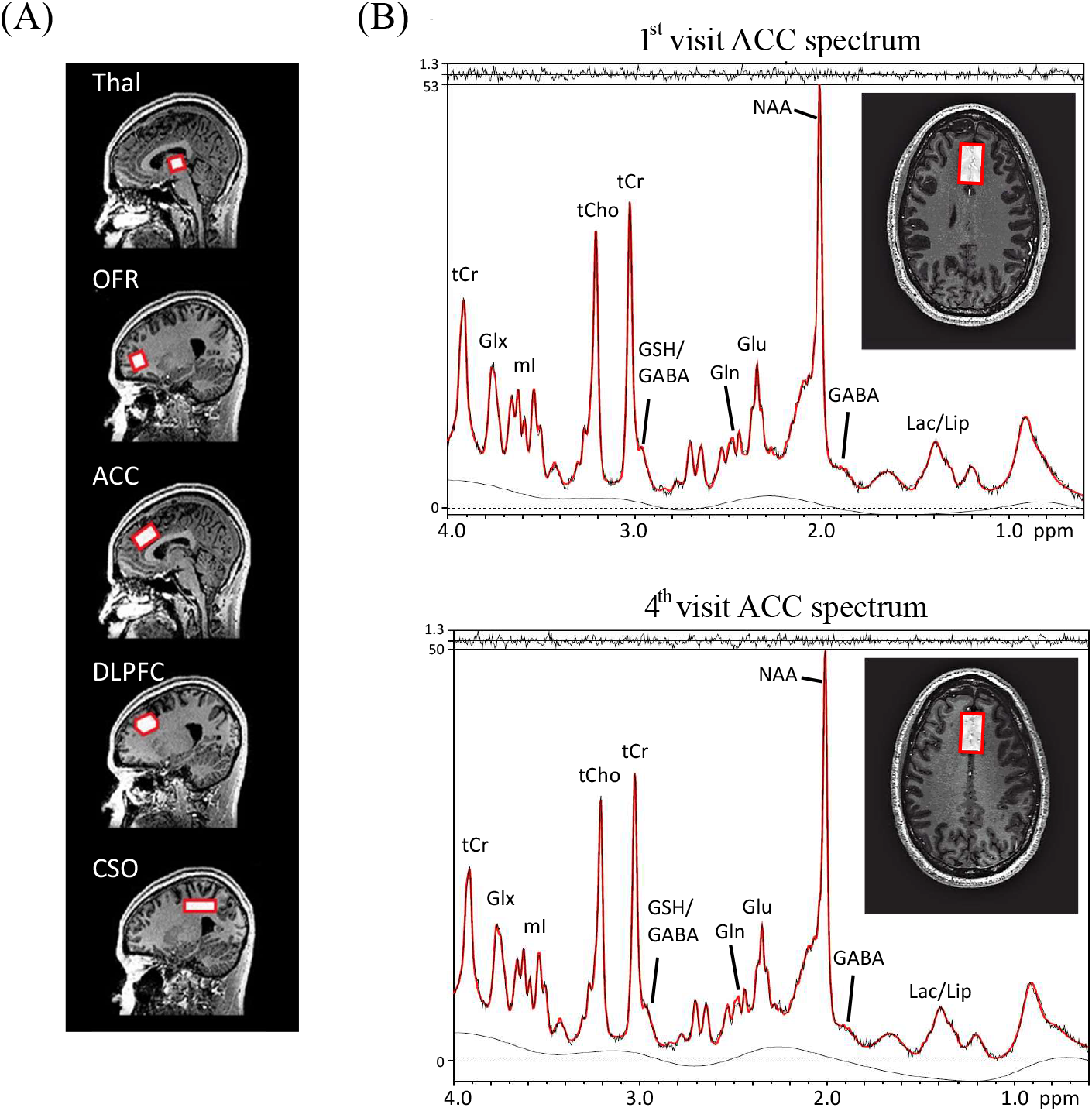
Magnetic Resonance Spectroscopy (MRS) Voxel Localizations and Representative Spectra. **(A)** Sagittal T_1_-weighted images showing the locations of the five brain regions (red boxes) used for MRS in this study. **(B)** ACC (anterior cingulate cortex) spectra recorded at the 1^st^ and 4^th^ visit in one subject, showing the voxel location overlaid on an axial T_1_-weighted image at both time points with results of the LCModel analysis; the LCModel output (red line) is superimposed on the original data (black line). The residual (fit - data) is shown at the top. Abbreviations: GABA indicates γ-aminobutyric acid; Gln, glutamine; Glu, glutamate; Glx, glutamate plus glutamine; GSH, glutathione; Lac, lactate; Lip, lipid; NAA, N-acetylaspartate; NAAG, N-acetylaspartyl glutamate; mI, myo-inositol; tCho, phosphocholine plus glycerophosphocholine; tCr, creatine plus phosphocreatine; Thal, thalamus; OFR, orbital frontal cortex; DLPFC, dorsolateral prefrontal cortex; and CSO, centrum semiovale.

### MR data processing

All spectra were analyzed with the LCModel software package (30) using a simulated basis set (31). Spectra were fitted between 0.2 and 4.0 ppm after eddy-current correction. Metabolite levels were estimated relative to the unsuppressed water signal. Full details of the spectral analysis methods have been described previously (12). MRS voxel composition was determined by segmenting the anatomical T_1_-weighted images into gray matter (GM), white matter (WM) and cerebrospinal fluid (CSF) using the SPM12 toolbox (32). Twenty metabolites were included in the LCModel basis set as described previously(12); after applying the previously described quality control criteria, the following metabolite concentration estimates were considered reliable enough for further statistical analysis: GABA, Glu, glutamine (Gln), GSH, lactate (Lac), myo-inositol (mI), NAA, N-acetyl aspartyl glutamate (NAAG), and total choline (tCho: phosphocholine plus glycerophosphocholine).

### Systematic bias between visits

To address whether there is systematic bias between visits, which could potentially affect our analysis results, for each subject we calculated the variance of 9 measured metabolite levels between visits. Our analysis showed that there were no uniform shifts in metabolite levels in any brain region (**Table S1-S5**), confirming that longitudinal changes in metabolites are unlikely caused by systematic bias.

### Return bias

To address return bias, which could potentially affect the analysis results, we compared the level of all measured metabolites in 5 brain regions obtained during the baseline visits for all study participants in our previous cross-sectional full cohort (baseline levels) (12) with those for the current longitudinal cohort. Linear regression with age, gender, race, and smoking status as covariates was performed in the HC only group, while linear regression with age, gender, race, smoking status, chlorpromazine (CPZ) equivalent dose, and duration of illness as covariates was performed in the FEP only group.

### Annual percentage change (APC)

To estimate rates of change of metabolite levels over time, annual percentage changes (APC) were calculated using the following expression:

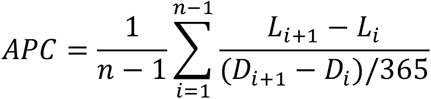

*L* is the level of metabolite, *D* is the date of the visit, *i* is the ith visit, *D_i,+1_* – *D_i_* is the number of days between two visits, and *n* is the total number of visits. Voxel tissue fractions were also investigated for changes over time using linear regression.

### Statistical analyses

To compare the APC between FEP patients and HC subjects, we performed the Bayesian two sample test and linear regression controlling for age, gender, race, and smoking status. The Benjamini-Hochberg (BH) procedure, a popular method for controlling the false discovery rate (FDR), was used for multiple comparison correction across all the measured metabolites and brain regions. P values corrected with the BH procedure are presented as q values. Bayes factor (BF), a statistical index not dependent on sample size and other extraneous factors (33), together with q values were used to evaluate the significance of analysis results. Similarly, to test if the mean APC within the HC or FEP group was different from zero, one sample t-test and Bayesian test were performed. The analyte was considered significant if its BF was larger than 10 and its q value was smaller than 0.05.

Furthermore, linear regression was performed to study the correlation between brain metabolite levels and other factors including age, smoking status, duration of illness, and CPZ dose obtained during the baseline visit. The absolute values of the t values were used to estimate the impact of these factors on brain metabolite levels.

## Results

### Longitudinal cohort

**Table 1** provides the demographic information of 38 FEP patients and 48 HC subjects in the present longitudinal cohort. Most study participants were African American, reflecting the composition of the local population. Additionally, FEP patients had a greater incidence of tobacco use than HC subjects. Differences in these demographic factors between FEP patients and HC subjects were adjusted in subsequent statistical analyses. In short, the demographic characteristics of our previous cross-sectional cohort (full cohort) (12) and those of our current longitudinal cohort look similar, however, we statistically made sure that there was no return bias in the longitudinal cohort. Between the full cohort and the present longitudinal cohort, we observed no significant differences in any metabolite in any brain region in both the HC group and the FEP group (**Tables S6, S7**).

**Table 1.** Demographics of study participants. Data collected during the subjects’ baseline visit is shown. T-test was performed to compare the age between first episode psychosis (FEP) patients and healthy control (HC) subjects. Fisher exact test was used to compare gender, race, and smoking status between FEP patients and HC subjects. Abbreviations: sd indicates standard deviation; Y, yes; and N, No.

| Characteristics | FEP (n=38) | HC (n=48) | p-value |
| --- | --- | --- | --- |
| Age (mean ± sd in years) | 22.53 ± 4.33 | 23.67 ± 3.31 | 0.18 |
| Gender (Male/Female) | 27/11 | 26/22 | 0.17 |
| Race (AA/Caucasian/Asian/Other) | 25/10/2/1 | 29/15/1/3 | 0.69 |
| Smoking status (Y/N) | 12/26 | 4/44 | 0.01 |
| CPZ (mean ± sd in mg) | 274.11 ± 211.67 | N/A | N/A |
| DOI (mean ± sd in month) | 15.26 ± 9.54 | N/A | N/A |

We then addressed longitudinal changes in brain metabolites in the present longitudinal cohort and employed the APC as a quantitative indicator (see Method section). By assessing the longitudinal changes in 9 measured metabolites in 5 brain regions, we observed several patterns as described below.

### Longitudinal reductions in ACC Glu in both FEP patients and HC subjects

We observed a negative mean APC for ACC Glu in both FEP patients and HC subjects, indicating that ACC Glu levels were reduced over time (**Figure 2A, Table 2**). In HC subjects, the mean APC of ACC Glu was −2.51%, whereas in FEP patients the mean APC was −5.03%. One sample t-test and Bayesian test supported the significance in reduction over time in ACC Glu in both FEP patients and HC subjects (BF > 10 and q-value < 0.05) (**Figure 2A**, **Table 2**). These data indicate that longitudinal changes exist even at physiological levels in young adulthood. The longitudinal reduction in Glu was only observed in the ACC, not in other brain regions.

**Figure 2.**
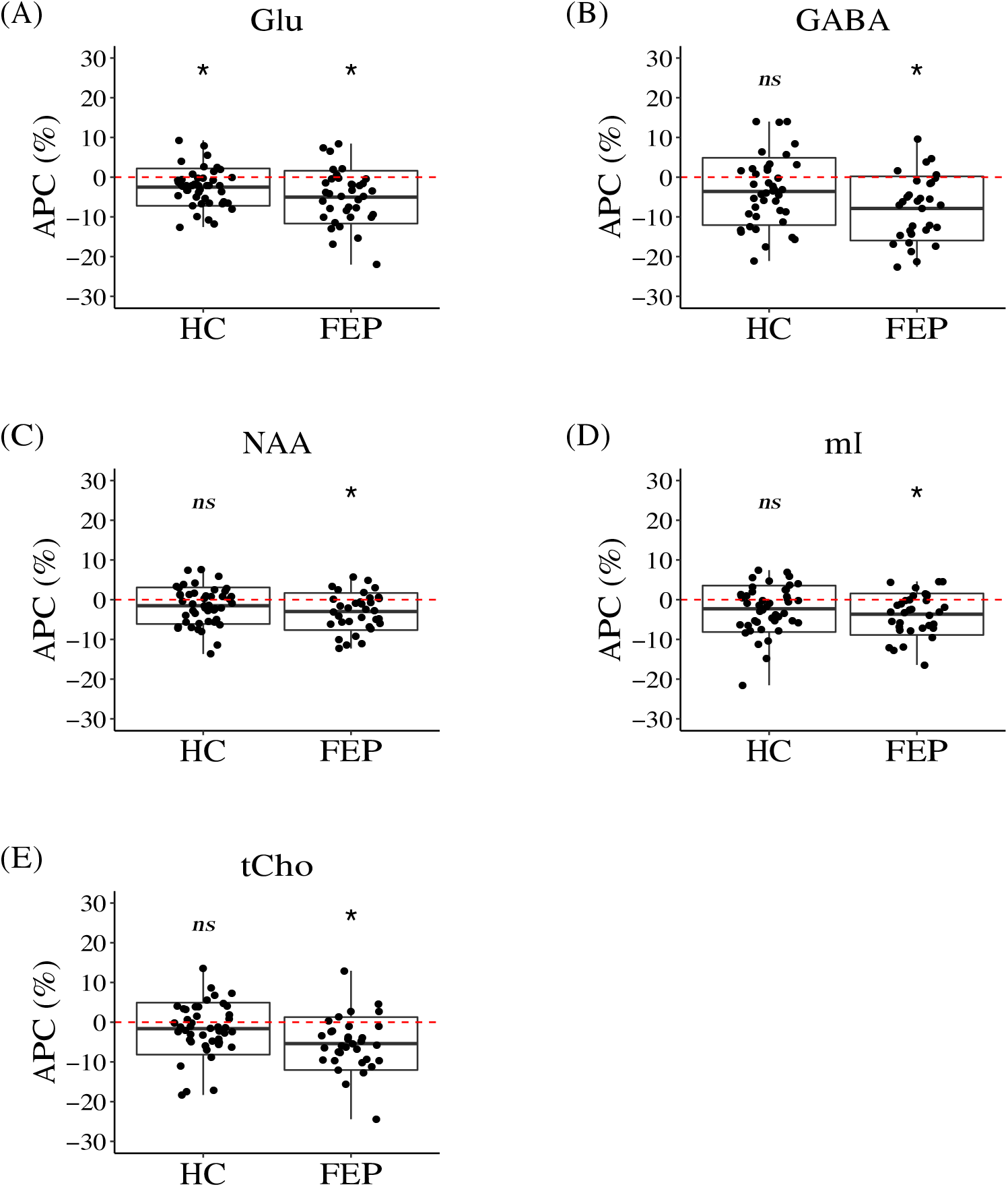
Boxplots of the Annual Percentage Change (APC) in Neurometabolites in the Anterior Cingulate Cortex (ACC). The red dotted line shows the value of zero. The box represents standard deviation and the solid line in the middle of the box shows the mean value of the APC. The black dots represent individual subjects. Symbol * denotes significant results, while symbol *ns* denotes results that didn’t reach the significant threshold (Bayes factor (BF) > 10 and q-value < 0.05). Abbreviations: Glu, glutamate; GABA, γ-aminobutyric acid; NAA, N-acetylaspartate; mI, myo-inositol; and tCho, phosphocholine plus glycerophosphocholine.

**Table 2.**
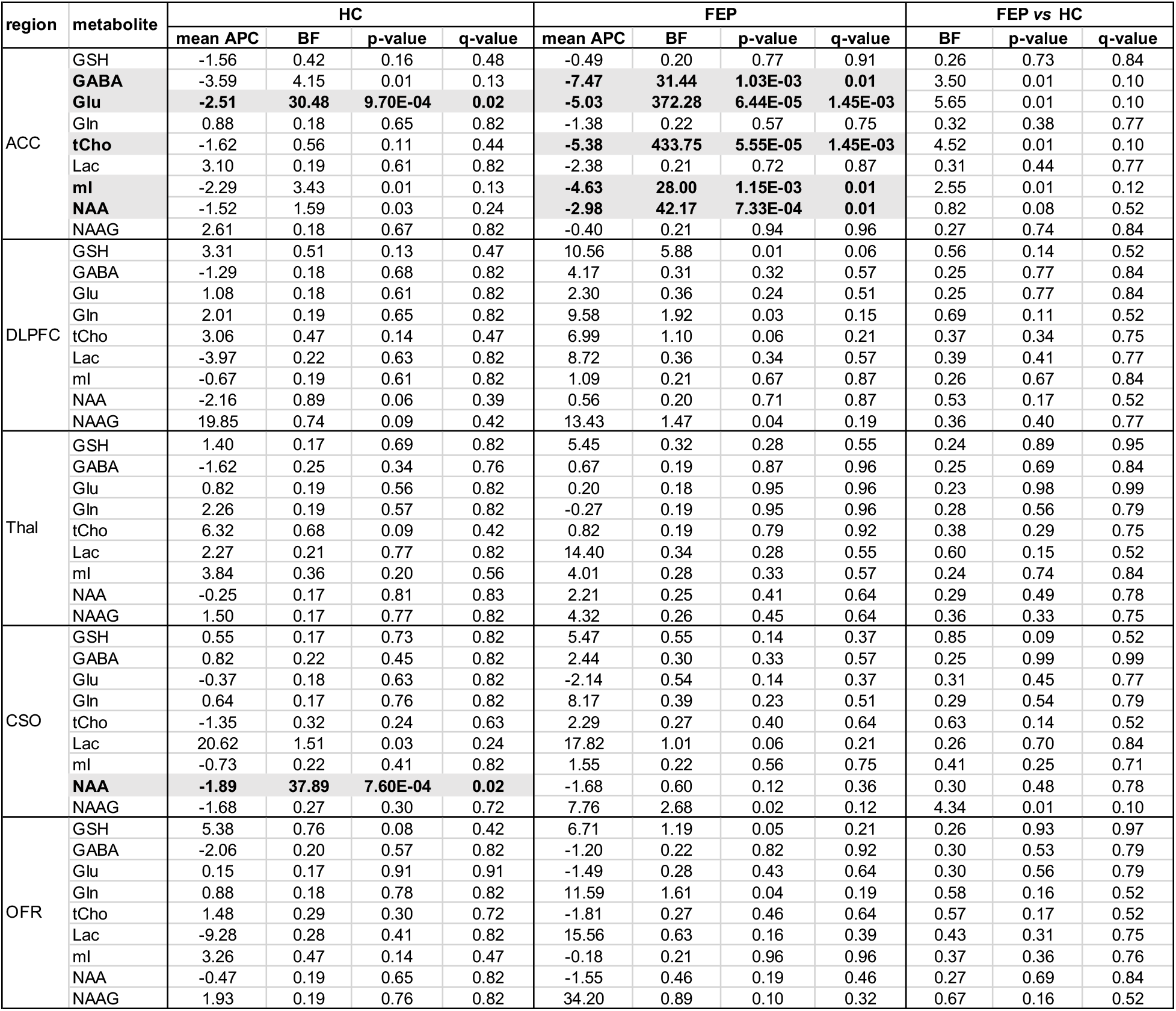
Longitudinal analysis of metabolites in the Anterior Cingulate Cortex (ACC). Mean annual percentage change (APC) was calculated to quantitively measure the longitudinal changes. A negative value indicates a decrease over time while a positive value indicates an increase. One sample Bayesian test and t-test were performed to check if the mean APC in one group (healthy control (HC) or first episode psychosis (FEP)) was significantly different from zero. Linear regression and Bayesian two sample test were performed to compare the APC between FEP patients and HC subjects. The Benjamini-Hochberg procedure was performed for multiple comparison correction. Significant results (Bayes factor (BF) > 10 and q-value < 0.05) are highlighted in bold with a gray shadow. Abbreviations: GABA indicates γ-aminobutyric acid; Gln, glutamine; Glu, glutamate; GSH, glutathione; Lac, lactate; NAA, N-acetylaspartate; NAAG, N-acetylaspartyl glutamate; mI, myo-inositol; tCho, phosphocholine plus glycerophosphocholine; ACC indicates anterior cingulate cortex; Thal, thalamus; DLPFC, dorsolateral prefrontal cortex; CSO, centrum semiovale; and OFR, orbital frontal region.

On average, FEP patients had a faster decline in ACC Glu compared to HC subjects, though it did not reach statistical significance (p-value = 0.01, q-value = 0.10, BF = 5.65). Note that, our previous study found that ACC Glu was significantly lower in FEP patients compared to HC subjects at baseline (12).

### Longitudinal reductions in GABA, NAA, mI, and tCho in the ACC in FEP patients

In addition to Glu, we observed significant longitudinal reductions in GABA, NAA, mI, and tCho in the ACC only in FEP patients (**Figure 2**, **Table 2**). The mean APC of GABA, NAA, mI, and tCho in FEP patients were −7.47%, −2.98%, −4.63%, and −5.38%, respectively. We didn’t observe significant differences in the APC between FEP patients and HC subjects. Note that at baseline, our previous study found that GABA and NAA in the ACC were significantly lower in FEP patients compared to HC subjects, while the difference in mI and tCho between FEP patients and HC subjects were not (12). These longitudinal reductions in GABA, NAA, mI, and tCho in FEP patients were only observed in the ACC, not in other brain regions.

In addition, we observed a significant reduction in CSO NAA in HC subjects (**Table 2**), but not in FEP patients. The longitudinal change in CSO NAA between FEP patients and HC subjects didn’t reach significance.

### FEP patients and HC subjects did not have significant longitudinal changes in GSH in any brain region

In the cross-sectional cohort we published for the 7T MRS study (12), the levels of GSH were significantly lower in the ACC and Thal in FEP patients compared with HC subjects. However, when we examined the longitudinal change in GSH, we didn’t observe a significant difference in the APC of GSH between FEP patients and HC subjects in any brain region (**Figure 3, Table 2**). In addition, the mean APC of GSH in every brain region in both FEP patients and HC subjects was not found to be significantly different from zero (**Figure 3, Table 2**). These results suggest that GSH levels are stable in 15- to 35-year-old subjects during a 4-year follow-up. There were several other metabolites that did not show longitudinal changes, but GSH was unique in having a significant difference at baseline between FEP patients and HC subjects without longitudinal changes.

**Figure 3.**
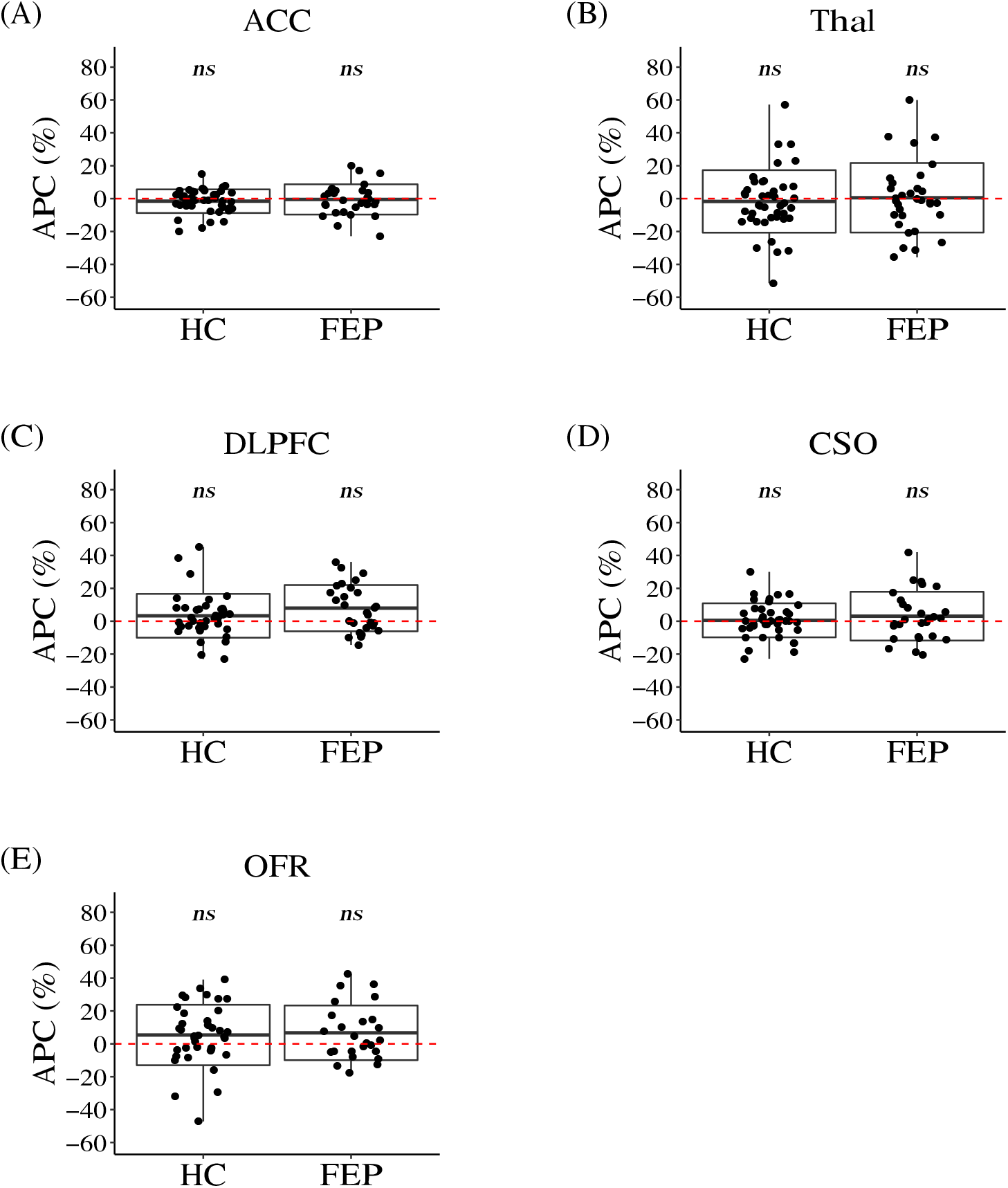
Boxplots of the Annual Percentage Change (APC) in Glutathione (GSH). The red dotted line shows the value of zero. The box represents standard deviation and the solid line in the middle of the box shows the mean value of the APC. The black dots represent individual subjects. Symbol * denotes significant results, while symbol *ns* denotes results that didn’t reach the significant threshold (Bayes factor (BF) > 10 and q-value < 0.05). Abbreviations: ACC, indicates anterior cingulate cortex; Thal, thalamus; DLPFC, dorsolateral prefrontal cortex; CSO, centrum semiovale; and OFR, orbital frontal cortex.

### Consideration of confounding factors that may affect the FEP group

In the above analyses, we considered age, gender, race, and smoking status as potential confounding factors for both FEP patients and HC subjects. In addition, the patient group may be influenced by intrinsic factors (disease-associated factors, such as speed and type of disease progression, as well as duration of illness (DOI)) and extrinsic factors (antipsychotic medication (CPZ dose)). To address this, we performed linear regression for metabolites with significant longitudinal changes (Glu, GABA, NAA, mI, and tCho in the ACC) in the FEP group. Since gender and race do not change over time, they were included in the linear regression as covariates. The severity and type of disease progression are difficult to include in the analysis, but DOI may reflect part of their impact. Together, we evaluated the effects of age, smoking status, antipsychotic medication, and DOI in FEP patients. We found that age had a significant impact on the level of ACC Glu and ACC GABA, while DOI had a significant impact on ACC NAA (**Figure S1**).

## Discussion

As far as we are aware, this study is the first to report longitudinal changes in brain metabolites in FEP patients and HC subjects in young adulthood using 7T MRS. There were three major findings in this study: (1) Glu showed a longitudinal decline only in the ACC in both FEP patients and HC subjects. (2) GABA, NAA, mI, and tCho showed a longitudinal decline only in the ACC in FEP patients, while NAA showed a longitudinal decline in the CSO in HC subjects; and (3) GSH levels in all the studied brain regions were relatively stable over time, although baseline levels of GSH in the ACC and Thal in FEP patients were lower than baseline levels in HC subjects(12). This data implies longitudinal changes in key brain metabolites both at the physiological and pathophysiological levels, which were preferentially represented in the ACC.

### Glu

Glu is the primary excitatory neurotransmitter in the adult human brain. The reduction of Glu in the pathological trajectory of patients with psychosis, including those with FEP, has been reported by multiple groups (7, 9, 11). Although there are reports that compared brain metabolite levels between young and aged populations (34), we believe that this study is the first to report that a longitudinal change in young adulthood occurs in healthy subjects. This implies dynamic changes in Glu even at the physiological level in people between 15 and 33 years old.

### GABA, NAA, mI, and tCho

GABA is the primary inhibitory neurotransmitter in the adult human brain. Some studies have shown that psychotic patients have lower levels of GABA compared to healthy controls (7, 11). NAA is the second most abundant metabolite in human brain (35). Its potential involvement in brain disorders such as SZ, Alzheimer’s disease, and brain injury has been reported (36). mI is utilized as a key element for intracellular signal transduction pathways (37). It is highly enriched in astrocytes and has been used as a marker of astrocyte activity (38). A meta-analysis reported a small, but significant, reduction in mI concentration in the medial frontal cortex in SZ (39).

Choline is an essential element for membrane synthesis and cholinergic neurotransmission. Interestingly, maternal prenatal choline deficiency has been strongly linked to subsequent development of SZ (40). Together, GABA, NAA, mI, and tCho has been reported to play important roles in neuropathology. Here, in this study, we observed significant longitudinal reductions in these metabolites in FEP patients, which could be a reflection of disease progress. Although any clinical study tends to stay descriptive, these observations may provide a useful hit to more mechanistic studies in animal models that mimic pathophysiology relevant to SZ and psychosis.

### GSH

One novel finding of the current study is that GSH has a near zero annual change in all brain regions. GSH is the most abundant non-enzymatic antioxidant in the central nervous system (41). Maintaining sufficient levels of GSH is important for protection against oxidative damage. MRS detection of GSH levels is usually presented as an indicator of antioxidant capacity in brain tissue. Our previous 7T MRS findings showed that there was a significantly lower GSH level in the ACC and thalamus in FEP patients compared to HC subjects (12). This suggested the presence of oxidative stress in the brain in early psychotic patients and provided evidence for the existence of a homeostatic imbalance in FEP patients (12, 42). In this study, we found that the GSH level did not have an obvious change longitudinally over a 4-year timespan for both FEP patients and HC subjects. Together, our findings suggest that GSH could be a trait marker for diagnostic applications. In addition, a related antioxidant or GSH-targeted treatment might be an alternative therapeutic pathway. Indeed, studies using animal models have suggested that redox imbalance and oxidative stress in adolescence later lead to cognitive and behavioral deficits relevant to psychotic disorders, and interventions in early stages can have a prophylactic impact (43, 44).

### Possible limitations

There are limitations in the current study, such as a limited number of brain regions examined and a narrow time window (4 year follow up in young adulthood). Future studies with a larger sample size, longer follow up, standardization of MRS techniques that could measure more brain regions, and unmedicated patients are encouraged to validate and expand the current findings.

### New analytic pipeline

In addition to novel findings in the longitudinal changes associated with physiological processes and pathological progressions, this study provides a straightforward analytic pipeline that leveraged the easy-to-understand concept of annual percentage change and well-developed statistical tests such as one sample test and linear regression. The analysis results are self-explanatory. Most importantly, this pipeline can be used in datasets with limited time points. This pipeline doesn’t depend on the type of dataset, which means that it can be applied not only to MRS data, but also to protein measurements, next generation sequencing data, clinical scales, or magnetic resonance imaging scans. Moreover, it can be carried out by researchers without statistical training.

## Conclusions

This study, for the first time, provided novel views of three longitudinal patterns in neurometabolites from both physiological and pathological viewpoints by leveraging the advantages of 7T MRS. GSH was found to have a near zero change in all 5 studied brain regions over time. Given that GSH levels were different in FEP patients and HC subjects at baseline, but stable over time, GSH may be a trait marker that could be used for diagnostic applications. Additionally, Glu, GABA, NAA, mI, and tCho in the ACC may be associated with the patient’s status and affected by physiological and pathological factors. These characteristics make them good state markers to study the mechanism of the development of psychiatric diseases.

## Acknowledgments

This study is supported by National Institutes of Mental Health Grants MH-092443 (to AS), MH-094268 (to AS), MH-105660 (to AS), and MH-107730 (to AS); foundation grants from Stanley (to AS), RUSK/S-R (to AS), and a NARSAD young investigator award from Brain and Behavior Research Foundation (to AS, KY). Study recruitment was in part funded by Mitsubishi Tanabe Pharm. Co. Ltd, Japan. The authors thank Yukiko Lema for suggestions for formatting the figures and her role in research management, and thank Dr. Mellisa A Landek-Salgado for scientific and English editions.

## References

1. Rae CD: A guide to the metabolic pathways and function of metabolites observed in human brain 1H magnetic resonance spectra. Neurochem Res 2014; 39:1–36

2. Poels EMP, Kegeles LS, Kantrowitz JT, et al.: Glutamatergic abnormalities in schizophrenia: a review of proton MRS findings. Schizophr Res 2014; 152:325–332

3. Stone JM, Dietrich C, Edden R, et al.: Ketamine effects on brain GABA and glutamate levels with 1H-MRS: relationship to ketamine-induced psychopathology. Mol Psychiatry 2012; 17:664–665

4. Mekle R, Mlynárik V, Gambarota G, et al.: MR spectroscopy of the human brain with enhanced signal intensity at ultrashort echo times on a clinical platform at 3T and 7T. Magn Reson Med 2009; 61:1279–1285

5. Pradhan S, Bonekamp S, Gillen JS, et al.: Comparison of single voxel brain MRS AT 3T and 7T using 32-channel head coils. Magn Reson Imaging 2015; 33:1013–1018

6. Tkác I, Oz G, Adriany G, et al.: In vivo 1H NMR spectroscopy of the human brain at high magnetic fields: metabolite quantification at 4T vs. 7T. Magn Reson Med 2009; 62:868–879

7. Reid MA, Salibi N, White DM, et al.: 7T Proton Magnetic Resonance Spectroscopy of the Anterior Cingulate Cortex in First-Episode Schizophrenia. Schizophr Bull 2019; 45:180–189

8. Brandt AS, Unschuld PG, Pradhan S, et al.: Age-related changes in anterior cingulate cortex glutamate in schizophrenia: A (1)H MRS Study at 7 Tesla. Schizophr Res 2016; 172:101–105

9. Kumar J, Liddle EB, Fernandes CC, et al.: Glutathione and glutamate in schizophrenia: a 7T MRS study. Mol Psychiatry 2018; 1–10

10. Rowland LM, Pradhan S, Korenic S, et al.: Elevated brain lactate in schizophrenia: a 7 T magnetic resonance spectroscopy study. Transl Psychiatry 2016; 6:e967

11. Thakkar KN, Rösler L, Wijnen JP, et al.: 7T Proton Magnetic Resonance Spectroscopy of Gamma-Aminobutyric Acid, Glutamate, and Glutamine Reveals Altered Concentrations in Patients With Schizophrenia and Healthy Siblings. Biol Psychiatry 2017; 81:525–535

12. Wang AM, Pradhan S, Coughlin JM, et al.: Assessing Brain Metabolism With 7-T Proton Magnetic Resonance Spectroscopy in Patients With First-Episode Psychosis. JAMA Psychiatry 2019; 76:314–323

13. Marsman A, Mandl RCW, Klomp DWJ, et al.: GABA and glutamate in schizophrenia: a 7 T ^1^H-MRS study. Neuroimage Clin 2014; 6:398–407

14. Merritt K, McGuire P, Egerton A: Relationship between Glutamate Dysfunction and Symptoms and Cognitive Function in Psychosis. Front Psychiatry 2013; 4:151

15. Merritt K, Perez-Iglesias R, Sendt K-V, et al.: Remission from antipsychotic treatment in first episode psychosis related to longitudinal changes in brain glutamate. NPJ Schizophr 2019; 5:12

16. Egerton A, Modinos G, Ferrera D, et al.: Neuroimaging studies of GABA in schizophrenia: a systematic review with meta-analysis. Transl Psychiatry 2017; 7:e1147

17. de la Fuente-Sandoval C, Reyes-Madrigal F, Mao X, et al.: Prefrontal and Striatal Gamma-Aminobutyric Acid Levels and the Effect of Antipsychotic Treatment in First-Episode Psychosis Patients. Biol Psychiatry 2018; 83:475–483

18. Marsman A, van den Heuvel MP, Klomp DWJ, et al.: Glutamate in schizophrenia: a focused review and meta-analysis of ^1^H-MRS studies. Schizophr Bull 2013; 39:120–129

19. Théberge J, Williamson KE, Aoyama N, et al.: Longitudinal grey-matter and glutamatergic losses in first-episode schizophrenia. Br J Psychiatry 2007; 191:325–334

20. Cadena EJ, White DM, Kraguljac NV, et al.: A Longitudinal Multimodal Neuroimaging Study to Examine Relationships Between Resting State Glutamate and Task Related BOLD Response in Schizophrenia. Front Psychiatry 2018; 9:632

21. Egerton A, Bhachu A, Merritt K, et al.: Effects of Antipsychotic Administration on Brain Glutamate in Schizophrenia: A Systematic Review of Longitudinal 1H-MRS Studies. Front Psychiatry 2017; 8:66

22. Egerton A, Broberg BV, Van Haren N, et al.: Response to initial antipsychotic treatment in first episode psychosis is related to anterior cingulate glutamate levels: a multicentre 1H-MRS study (OPTiMiSE). Mol Psychiatry 2018; 23:2145–2155

23. Egerton A, Stone JM, Chaddock CA, et al.: Relationship between brain glutamate levels and clinical outcome in individuals at ultra high risk of psychosis. Neuropsychopharmacology 2014; 39:2891–2899

24. Zeger SL, Liang KY: An overview of methods for the analysis of longitudinal data. Stat Med 1992;11:1825–1839

25. Younes L, Albert M, Moghekar A, et al.: Identifying Changepoints in Biomarkers During the Preclinical Phase of Alzheimer’s Disease. Front Aging Neurosci 2019; 11:74

26. Caruana EJ, Roman M, Hernández-Sánchez J, et al.: Longitudinal studies. J Thorac Dis 2015; 7:E537–E540

27. Garcia TP, Marder K: Statistical Approaches to Longitudinal Data Analysis in Neurodegenerative Diseases: Huntington’s Disease as a Model. Curr Neurol Neurosci Rep 2017; 17:14

28. Gruetter R: Automatic, localized in vivo adjustment of all first- and second-order shim coils. Magn Reson Med 1993; 29:804–811

29. Versluis MJ, Kan HE, van Buchem MA, et al.: Improved signal to noise in proton spectroscopy of the human calf muscle at 7 T using localized B1 calibration. Magn Reson Med 2010; 63:207–211

30. Provencher SW: Estimation of metabolite concentrations from localized in vivo proton NMR spectra. Magn Reson Med 1993; 30:672–679

31. Soher B, Semanchuk P, Todd D, et al.: Vespa: Integrated applications for RF pulse design, spectral simulation and MRS data analysis. Proc. Intl. Soc. Mag. Reson. Med. 2011.

32. Eickhoff SB, Stephan KE, Mohlberg H, et al.: A new SPM toolbox for combining probabilistic cytoarchitectonic maps and functional imaging data. Neuroimage 2005; 25:1325–1335

33. Jarosz A, Wiley J: What Are the Odds? A Practical Guide to Computing and Reporting Bayes Factors. J Probl Solving 2014; 7:2

34. Cleeland C, Pipingas A, Scholey A, et al.: Neurochemical changes in the aging brain: A systematic review. Neurosci Biobehav Rev 2019; 98:306–319

35. Miyake M, Kakimoto Y, Sorimachi M: A gas chromatographic method for the determination of N-acetyl-L-aspartic acid, N-acetyl-alpha-aspartylglutamic acid and beta-citryl-L-glutamic acid and their distributions in the brain and other organs of various species of animals. J Neurochem 1981; 36:804–810

36. Moffett JR, Ross B, Arun P, et al.: N-Acetylaspartate in the CNS: From Neurodiagnostics to Neurobiology. Prog Neurobiol 2007; 81:89–131

37. Gillaspy GE: The cellular language of myo-inositol signaling. New Phytologist 2011; 192:823–839

38. Harris JL, Choi I-Y, Brooks WM: Probing astrocyte metabolism in vivo: proton magnetic resonance spectroscopy in the injured and aging brain. Front Aging Neurosci 2015; 7:202

39. Das TK, Dey A, Sabesan P, et al.: Putative Astroglial Dysfunction in Schizophrenia: A Meta-Analysis of 1H-MRS Studies of Medial Prefrontal Myo-Inositol. Front Psychiatry 2018; 9:438

40. Freedman R, Ross RG: Prenatal choline and the development of schizophrenia. Shanghai Arch Psychiatry 2015; 27:90–102

41. Ramírez-Expósito MJ, Martínez-Martos JM: The Delicate Equilibrium between Oxidants and Antioxidants in Brain Glioma. Curr Neuropharmacol 2019; 17:342–351

42. Landek-Salgado MA, Faust TE, Sawa A: Molecular substrates of schizophrenia: homeostatic signaling to connectivity. Mol Psychiatry 2016; 21:10–28

43. Cabungcal J-H, Counotte DS, Lewis EM, et al.: Juvenile Antioxidant Treatment Prevents Adult Deficits in a Developmental Model of Schizophrenia. Neuron 2014; 83:1073–1084

44. Sawa A, Seidman LJ: Is Prophylactic Psychiatry around the Corner? Combating Adolescent Oxidative Stress for Adult Psychosis and Schizophrenia. Neuron 2014; 83:991–993

