## Supplementary Document for "Longitudinal changes in brain metabolites in healthy subjects and patients with first episode psychosis (FEP): a 7-Tesla MRS study"

**Table S1**. **Analysis results of systematic bias between visits in anterior cingulate cortex.**

Variations between the first and second visits were calculated. A negative variation means that there is a decrease in the second visit and is highlighted in red. A positive variation means that there is an increase in the second visit and is highlighted in green. A variation of "0.00" means that there is no change between visits and is highlighted in gray. There was not a systematic increase or decrease observed between visits. This suggests that there is not a technique batch effect that may affect the longitudinal analysis results. An empty cell means the data was excluded from analysis due to low quality. Abbreviations: GABA indicates γ-aminobutyric acid; Gln, glutamine; Glu, glutamate; GSH, glutathione; Lac, lactate; NAA, N-acetylaspartate; NAAG, N-acetylaspartyl glutamate; mI, myo-inositol; tCho, phosphocholine plus glycerophosphocholine.

| Subject | GSH | GABA | Glu | Gln | tCho | Lac | mI | NAA | NAAG |
|---|---|---|---|---|---|---|---|---|---|
| 1001 | -0.26 | -0.41 | -1.87 | 0.59 | -0.18 | -0.07 | -1.20 | -1.21 | 0.03 |
| 1005 | 0.23 | 0.09 | 0.59 | -0.13 | -0.04 | -0.06 | -0.21 | 0.25 | -0.45 |
| 1006 | -0.03 | -0.22 | -0.52 | -0.17 | -0.16 | -0.21 | -0.39 | 0.23 | -0.30 |
| 1007 | 0.18 | 0.02 | -0.09 | 0.14 | -0.10 | -0.01 | 0.16 | 0.13 | -0.19 |
| 1011 | -0.16 | -0.09 | -0.02 | 0.03 | 0.06 | 0.00 | -0.05 | 0.02 | -0.10 |
| 1012 | 0.05 | 0.16 | 0.49 | -0.02 | 0.17 | 0.21 | 0.19 | -0.03 | 0.20 |
| 1014 | 0.08 | -0.15 | -0.15 | -0.01 | -0.07 | -0.28 | -0.33 | 0.04 | |
| 1018 | 0.15 | -0.20 | 0.11 | -0.05 | 0.01 | -0.02 | -0.14 | 0.36 | 0.05 |
| 1022 | 0.01 | 0.05 | 0.30 | 0.26 | 0.04 | 0.15 | 0.29 | 0.42 | 0.03 |
| 1026 | 0.13 | -0.16 | -0.07 | -0.16 | -0.06 | | 0.10 | 0.47 | |
| 1033 | -0.19 | -0.18 | -0.41 | -0.17 | -0.08 | | -0.47 | -0.29 | 0.23 |
| 1038 | 0.18 | -0.22 | -1.33 | -0.26 | -0.11 | 0.55 | -1.92 | 0.31 | |
| 1039 | -0.28 | -0.20 | -0.36 | 0.66 | -0.14 | -0.09 | -0.80 | -0.13 | -0.47 |
| 1040 | -0.07 | -0.33 | -0.72 | -0.30 | -0.16 | -0.14 | -0.36 | -0.53 | |
| 1042 | 0.17 | 0.08 | 0.65 | 0.43 | -0.07 | | 0.00 | 0.26 | 0.09 |
| 1044 | 0.13 | -0.62 | -1.14 | | -0.32 | 0.07 | -1.26 | -0.37 | -0.52 |
| 1047 | | 0.53 | 0.66 | 0.16 | -0.22 | 0.35 | -0.13 | 0.41 | -0.14 |
| 1050 | 0.15 | 0.29 | 0.26 | 0.20 | -0.07 | -0.04 | -0.12 | 0.20 | -0.27 |
| 1051 | -0.05 | -0.34 | -0.35 | -0.25 | -0.15 | 0.16 | 0.16 | -0.26 | -0.12 |
| 1052 | 0.14 | -0.06 | 0.16 | -0.03 | 0.01 | 0.15 | 0.35 | 0.46 | -0.12 |
| 1055 | -0.05 | | -1.27 | | -0.12 | -0.21 | 0.32 | -0.37 | |
| 1056 | 0.07 | -0.08 | -0.66 | -0.29 | -0.02 | 0.02 | -0.14 | 0.22 | -0.22 |
| 1069 | -0.15 | -0.34 | -0.26 | | -0.33 | -0.09 | -0.60 | -0.31 | -0.06 |
| 1070 | -0.05 | -0.16 | -0.30 | 0.19 | -0.20 | -0.06 | 0.07 | -0.42 | |
| 1082 | -0.02 | -0.58 | -0.78 | -0.39 | 0.05 | -0.11 | -0.50 | -0.28 | |
| 1085 | 0.27 | -0.19 | -0.78 | -0.14 | | -0.24 | 0.21 | -0.07 | -0.11 |
| 1090 | | -0.34 | -0.39 | 0.10 | | -0.05 | -0.20 | -0.69 | -0.11 |
| 1091 | | -0.18 | -0.75 | -0.53 | -0.13 | -0.15 | -0.01 | -0.11 | |
| 1105 | 0.00 | -0.23 | -0.95 | -0.04 | -0.14 | | | | |

(continued on next page)

| | | | | | | | | | |
|---|---|---|---|---|---|---|---|---|---|
| 1109 | | -0.02 | -1.27 | -0.48 | -0.13 | | -0.01 | -0.80 | |
| 1118 | -0.19 | -0.08 | -0.70 | -0.06 | | -0.15 | -0.42 | -1.00 | 0.24 |
| 1119 | 0.21 | -0.01 | -0.09 | 0.42 | -0.04 | 0.14 | -0.08 | -0.34 | 0.06 |
| 1122 | -0.38 | | -1.11 | 0.39 | -0.11 | -0.41 | -0.54 | -0.76 | -0.01 |
| 1132 | -0.02 | -0.61 | -1.58 | -0.50 | -0.20 | -0.12 | -0.53 | | |
| 1136 | | | -0.65 | 0.01 | -0.09 | | -0.40 | -0.21 | -0.12 |
| 2001 | 0.14 | -0.08 | -0.31 | -0.10 | 0.06 | -0.18 | -0.04 | 0.06 | 0.01 |
| 2002 | -0.07 | -0.06 | -0.54 | 0.20 | 0.00 | 0.18 | -0.26 | -0.42 | -0.12 |
| 2003 | -0.07 | -0.07 | -0.52 | -0.16 | -0.04 | -0.13 | 0.14 | 0.22 | 0.19 |
| 2005 | 0.11 | -0.13 | -0.47 | -0.12 | 0.02 | 0.11 | -0.06 | -0.06 | 0.11 |
| 2007 | 0.18 | 0.51 | 1.59 | 0.85 | 0.23 | -0.13 | 0.63 | 1.20 | 0.27 |
| 2009 | 0.05 | 0.01 | -0.46 | -0.51 | 0.05 | 0.04 | -0.19 | -0.21 | -0.07 |
| 2010 | -0.25 | 0.12 | -0.20 | -0.41 | 0.06 | 0.02 | -0.47 | 0.11 | 0.16 |
| 2014 | 0.03 | 0.14 | 0.49 | 0.10 | 0.06 | 0.08 | 0.05 | 0.06 | 0.07 |
| 2016 | -0.04 | 0.14 | 0.20 | 0.54 | -0.09 | 0.03 | -0.18 | -0.34 | -0.07 |
| 2019 | 0.06 | 0.01 | -0.06 | 0.35 | 0.14 | 0.00 | 0.52 | -0.29 | -0.13 |
| 2022 | 0.06 | -0.22 | -0.48 | 0.07 | 0.00 | -0.07 | -0.26 | -0.25 | -0.23 |
| 2023 | 0.06 | 0.11 | -0.49 | 0.20 | -0.08 | -0.01 | -0.31 | -0.19 | 0.18 |
| 2024 | -0.12 | -0.08 | 0.40 | 0.12 | -0.06 | 0.17 | 0.02 | 0.04 | 0.09 |
| 2026 | 0.20 | -0.11 | -0.83 | -0.42 | -0.05 | -0.22 | 0.38 | 0.02 | -0.31 |
| 2028 | 0.12 | -0.17 | 0.18 | 0.21 | -0.07 | 0.00 | -0.21 | 0.12 | -0.10 |
| 2029 | 0.00 | -0.03 | 0.23 | -0.11 | -0.15 | 0.09 | | 0.07 | 0.08 |
| 2032 | -0.09 | -0.02 | 0.61 | 0.10 | 0.12 | -0.10 | 0.41 | 0.30 | 0.05 |
| 2038 | -0.01 | 0.24 | -0.23 | -0.17 | -0.04 | -0.11 | 0.25 | 0.13 | -0.11 |
| 2041 | -0.23 | 0.03 | -1.06 | -0.77 | -0.29 | 0.04 | -0.76 | -0.24 | -0.01 |
| 2042 | -0.14 | -0.07 | -0.43 | 0.12 | -0.04 | -0.12 | -0.21 | 0.03 | -0.16 |
| 2047 | -0.03 | 0.06 | -0.13 | 0.08 | 0.01 | 0.01 | 0.02 | 0.08 | -0.25 |
| 2048 | -0.10 | | -0.39 | -0.38 | -0.04 | -0.18 | -0.47 | -0.45 | |
| 2052 | | 0.03 | 0.31 | 0.11 | -0.22 | 0.01 | 0.20 | -0.11 | 0.31 |
| 2053 | -0.05 | -0.09 | 0.06 | 0.19 | | -0.02 | -0.25 | -0.51 | 0.16 |
| 2056 | 0.05 | | 0.28 | 0.10 | 0.16 | | 0.23 | -0.06 | |
| 2057 | 0.06 | | -0.18 | -0.04 | -0.05 | 0.40 | -0.01 | 0.40 | -0.32 |
| 2061 | | -0.08 | -0.59 | 0.15 | 0.05 | | 0.31 | 0.02 | -0.07 |
| 2062 | -0.21 | -0.12 | -0.99 | -0.13 | -0.27 | -0.08 | -1.25 | -0.56 | 0.55 |
| 2066 | 0.01 | -0.12 | -0.31 | 0.01 | -0.10 | -0.22 | -0.32 | -0.44 | 0.07 |
| 2067 | 0.05 | 0.23 | | -0.12 | 0.07 | 0.04 | -0.33 | 0.54 | -0.30 |
| 2068 | -0.28 | -0.34 | -0.85 | -0.23 | -0.02 | -0.22 | -0.34 | -0.89 | -0.05 |
| 2069 | -0.11 | 0.39 | -0.34 | 0.07 | -0.17 | -0.10 | -0.76 | -0.33 | 0.14 |
| 2070 | -0.02 | -0.28 | -0.52 | -0.14 | 0.08 | -0.10 | 0.45 | -0.35 | 0.03 |
| 2071 | -0.02 | 0.24 | -0.32 | 0.17 | 0.03 | -0.45 | | 0.23 | 0.18 |
| 2073 | 0.20 | -0.10 | -0.05 | 0.29 | 0.10 | -0.13 | 0.13 | 0.26 | -0.24 |
| 2075 | -0.02 | | -0.27 | -0.17 | -0.03 | 0.04 | 0.24 | -0.02 | -0.15 |
| 2078 | -0.07 | | -0.33 | -0.35 | -0.27 | 0.16 | -0.05 | 0.21 | -0.16 |
| 2080 | 0.14 | -0.10 | -0.04 | 0.21 | 0.00 | 0.01 | 0.07 | -0.29 | -0.02 |
| 2081 | -0.02 | 0.33 | -0.18 | 0.02 | -0.05 | 0.12 | -0.32 | 0.06 | 0.02 |
| 2082 | | -0.52 | -0.02 | | -0.22 | -0.03 | 0.11 | 0.06 | 0.22 |
| 2083 | 0.11 | -0.17 | | | -0.02 | 0.23 | -0.26 | 0.20 | -0.22 |
| 2084 | -0.08 | 0.00 | -0.05 | | -0.09 | 0.13 | -0.09 | 0.14 | -0.16 |
| 2087 | -0.16 | -0.36 | -0.99 | -0.47 | | 0.00 | -0.64 | -0.57 | -0.24 |
| 2099 | | -0.32 | -0.56 | -0.35 | -0.04 | -0.09 | -0.39 | -0.44 | -0.31 |
| 2101 | 0.09 | -0.28 | | 0.48 | -0.07 | -0.08 | -0.30 | -0.66 | 0.10 |
| 2102 | -0.36 | -0.16 | -0.70 | -0.13 | -0.18 | -0.25 | -0.61 | -0.92 | 0.05 |
| 2111 | -0.01 | -0.21 | -0.21 | 0.04 | -0.02 | | -0.26 | -0.50 | 0.11 |
| 2112 | -0.25 | -0.20 | -0.77 | | | -0.05 | -0.36 | -0.94 | -0.09 |

**Table S2**. **Analysis results of systematic bias between visits in thalamus.**

Variations between the first and second visits were calculated. A negative variation means that there is a decrease in the second visit and is highlighted in red. A positive variation means that there is an increase in the second visit and is highlighted in green. A variation of "0.00" means that there is no change between visits and is highlighted in gray. There was not a systematic increase or decrease observed between visits. This suggests that there is not a technique batch effect that may affect the longitudinal analysis results. An empty cell means the data was excluded from analysis due to low quality. Abbreviations: GABA indicates γ-aminobutyric acid; Gln, glutamine; Glu, glutamate; GSH, glutathione; Lac, lactate; NAA, N-acetylaspartate; NAAG, N-acetylaspartyl glutamate; mI, myo-inositol; tCho, phosphocholine plus glycerophosphocholine.

| Subject | GSH | GABA | Glu | Gln | tCho | Lac | mI | NAA | NAAG |
|---|---|---|---|---|---|---|---|---|---|
| 1001 | -0.27 | -0.08 | -1.50 | 0.64 | -0.30 | | -1.08 | -1.02 | 0.00 |
| 1005 | 0.48 | -0.63 | 0.29 | 0.09 | 0.28 | -0.26 | 0.86 | 0.20 | 0.33 |
| 1006 | 0.75 | 0.90 | 2.88 | 0.91 | 0.63 | -0.06 | 1.82 | 3.13 | |
| 1007 | -0.12 | 0.05 | -0.28 | -0.01 | -0.06 | -0.09 | -0.44 | 0.11 | -0.03 |
| 1011 | 0.04 | -0.28 | -0.07 | -0.12 | 0.06 | 0.09 | 0.25 | 0.14 | -0.17 |
| 1012 | 0.65 | 0.33 | 1.34 | | 0.26 | -0.31 | -0.02 | 0.89 | -0.12 |
| 1014 | | 0.07 | -0.64 | -0.34 | -0.22 | 0.36 | -1.03 | -0.30 | -0.19 |
| 1022 | 0.80 | 0.02 | 0.93 | -0.30 | -0.11 | -0.45 | 1.61 | 0.26 | 0.27 |
| 1026 | -0.03 | 0.05 | 0.34 | -0.04 | 0.31 | 0.22 | 1.02 | 0.45 | 0.44 |
| 1031 | -0.63 | -0.58 | -1.57 | | -0.53 | | -0.68 | -1.29 | |
| 1032 | 0.60 | -0.23 | -0.17 | 0.82 | -0.14 | 0.08 | -1.37 | 0.53 | -0.17 |
| 1033 | -0.03 | 0.22 | 0.34 | -0.09 | -0.01 | -0.05 | -0.15 | -0.06 | -0.06 |
| 1038 | -0.09 | -0.26 | -0.06 | -0.17 | -0.51 | 0.37 | -1.77 | -0.65 | 0.43 |
| 1039 | -0.76 | -0.72 | -1.43 | | -0.38 | | -0.75 | -1.31 | 1.22 |
| 1040 | -0.83 | -0.49 | -0.57 | -0.24 | -0.34 | | 0.05 | -0.28 | |
| 1042 | 0.37 | -0.21 | 0.76 | -0.71 | 0.06 | 0.46 | 0.38 | 1.49 | |
| 1044 | -0.10 | | 0.07 | 0.06 | 0.05 | -0.28 | -0.20 | -0.77 | -0.04 |
| 1047 | 0.56 | 0.62 | -0.17 | | -0.10 | | 0.52 | 0.90 | -1.13 |
| 1050 | 0.31 | 0.50 | 0.70 | -0.40 | 0.29 | -0.32 | 0.92 | 1.49 | -0.51 |
| 1051 | 0.48 | 0.33 | -0.20 | -0.08 | 0.23 | 0.34 | 1.45 | -0.04 | 0.29 |
| 1052 | -0.24 | 0.51 | 0.07 | 0.33 | -0.07 | 0.08 | -0.18 | -0.88 | 0.43 |
| 1055 | -0.14 | -0.95 | -0.79 | | | 0.95 | -1.07 | 0.12 | 0.57 |
| 1056 | -0.21 | 0.00 | -0.65 | -0.27 | -0.16 | 0.63 | -0.25 | -0.10 | 0.08 |
| 1070 | 0.87 | 1.04 | 2.61 | | 0.67 | -0.58 | 1.15 | 2.94 | 0.35 |
| 1082 | | -0.88 | -0.42 | -0.41 | 0.06 | | 2.61 | 0.92 | -0.30 |
| 1085 | -0.67 | | -0.63 | -0.16 | 0.08 | -0.03 | 0.89 | -0.06 | -0.21 |
| 1090 | 0.01 | -0.19 | -1.07 | 0.12 | 0.08 | -0.18 | -0.71 | -0.93 | 0.06 |
| 1091 | -0.05 | 0.59 | 0.18 | -0.44 | 0.39 | | 1.41 | 0.07 | 0.29 |
| 1105 | -0.18 | -1.12 | -2.38 | -1.06 | -0.38 | 0.20 | -1.95 | -0.79 | |

(continued on next page)

| | | | | | | | | | |
|---|---|---|---|---|---|---|---|---|---|
| 1109 | -0.52 | -1.14 | -1.94 | 0.32 | | -0.29 | -0.33 | | -0.54 |
| 1118 | 0.23 | 0.47 | 0.91 | -0.31 | 0.43 | 0.22 | 1.70 | 0.02 | 0.98 |
| 1119 | 0.00 | -0.46 | | 0.25 | -0.05 | -0.05 | 0.14 | 0.74 | |
| 1122 | -0.23 | -0.72 | 0.81 | 0.30 | -0.24 | -0.29 | -0.94 | -0.62 | -0.73 |
| 1132 | 0.15 | -0.01 | -1.48 | -0.18 | | -0.22 | -1.01 | -1.77 | -0.02 |
| 1136 | -0.36 | 0.12 | 0.12 | -0.18 | -0.29 | -0.20 | -0.43 | 0.32 | -0.37 |
| 2001 | 0.40 | 0.30 | 0.46 | 0.48 | -0.39 | | 1.28 | -0.08 | 0.40 |
| 2002 | 0.75 | -0.42 | 0.40 | 0.56 | 0.26 | -0.09 | -0.52 | 0.74 | -0.41 |
| 2003 | -0.53 | -0.11 | -0.96 | -0.25 | -0.12 | 0.02 | -0.23 | -0.22 | -0.56 |
| 2005 | 0.48 | -0.29 | 0.19 | 0.44 | 0.00 | 0.11 | 0.89 | 0.35 | 0.25 |
| 2007 | 0.07 | 0.25 | 0.38 | 0.32 | 0.14 | 0.00 | -0.20 | 0.45 | 0.11 |
| 2009 | 0.58 | -0.67 | -0.59 | 0.14 | -0.17 | 0.04 | 0.45 | 0.72 | -1.11 |
| 2010 | -0.21 | 0.17 | 0.05 | -0.27 | 0.01 | 0.08 | -0.54 | -0.13 | 0.19 |
| 2014 | -0.21 | 0.02 | -0.07 | -0.48 | -0.06 | | 1.23 | 0.00 | 0.23 |
| 2016 | -0.26 | -0.14 | -0.06 | 0.48 | -0.05 | | 0.33 | -0.11 | 0.00 |
| 2019 | 0.42 | 0.06 | 0.15 | 0.41 | 0.63 | 0.15 | 0.07 | 0.29 | 0.27 |
| 2022 | 0.19 | | 0.72 | 0.23 | -0.07 | 0.00 | 1.62 | 0.00 | 0.51 |
| 2023 | 0.54 | 0.14 | -0.61 | 0.04 | -0.19 | -0.27 | -1.56 | -0.03 | -0.29 |
| 2024 | -0.33 | -0.05 | -0.49 | -0.64 | -0.17 | -0.04 | -0.52 | 0.21 | 0.04 |
| 2026 | -0.13 | -0.81 | -1.40 | -0.41 | -0.02 | | -0.63 | -1.07 | -0.53 |
| 2028 | 0.00 | -0.04 | -1.03 | 0.15 | -0.15 | -0.16 | -0.67 | -0.94 | -0.33 |
| 2029 | -0.25 | -0.29 | 0.58 | -0.20 | -0.22 | 0.02 | -1.59 | -0.04 | 0.08 |
| 2032 | -0.96 | -0.22 | -0.83 | -0.30 | -0.07 | | 0.00 | -1.15 | 0.42 |
| 2038 | -0.04 | 0.54 | 0.36 | 0.37 | 0.09 | | -0.08 | 0.06 | 0.04 |
| 2041 | 0.19 | -0.10 | 0.32 | -0.76 | -0.04 | | 0.06 | 0.95 | -0.56 |
| 2042 | -0.23 | 0.43 | -0.73 | -0.11 | 0.38 | | 0.00 | -0.30 | 0.34 |
| 2047 | -1.13 | -0.23 | -0.90 | | 0.60 | -0.60 | 0.52 | -1.09 | -0.01 |
| 2048 | 0.34 | 0.19 | 0.66 | 0.33 | 0.40 | 0.08 | 0.51 | 0.38 | 0.30 |
| 2050 | -0.22 | -0.42 | -1.13 | 0.07 | -0.08 | -0.13 | 0.03 | -1.55 | 0.65 |
| 2052 | 0.31 | 0.04 | 1.78 | 0.43 | -0.19 | -0.10 | 0.66 | -0.16 | -0.07 |
| 2053 | 0.58 | 0.41 | 1.36 | 0.70 | 0.19 | | -0.20 | 1.03 | 0.30 |
| 2056 | 0.02 | 0.10 | 0.30 | 0.27 | 0.12 | -0.24 | 0.36 | -0.43 | |
| 2057 | 0.35 | 0.35 | 0.76 | | 0.86 | 0.26 | 1.74 | 0.50 | -1.32 |
| 2066 | -0.11 | 0.23 | 1.40 | -0.65 | 0.19 | | 0.36 | 1.38 | |
| 2067 | 0.13 | 0.01 | -0.25 | -0.46 | 0.23 | | -1.85 | 0.04 | 0.05 |
| 2068 | -0.02 | -0.24 | -0.77 | 0.00 | -0.11 | -0.27 | -0.02 | -0.01 | -0.16 |
| 2069 | 0.08 | 0.10 | -0.06 | -0.16 | -0.24 | 0.24 | -0.63 | 0.60 | -1.13 |
| 2070 | -0.07 | 0.02 | -0.69 | -0.08 | -0.14 | -0.47 | -0.10 | -0.85 | 0.03 |
| 2071 | 0.06 | 0.22 | -0.57 | 0.73 | 0.06 | | 2.63 | -0.32 | 0.33 |
| 2073 | -0.22 | 0.24 | -0.38 | -0.57 | 0.02 | 0.03 | 0.72 | -0.40 | 0.34 |
| 2075 | -0.26 | -0.54 | -0.27 | -0.35 | 0.05 | | 0.22 | -0.34 | -0.63 |
| 2078 | -0.19 | -0.24 | 0.65 | -0.29 | 0.14 | 0.17 | -0.06 | 0.40 | -0.37 |
| 2080 | -0.12 | -0.24 | 0.22 | -0.02 | | | -0.53 | -0.07 | -0.34 |
| 2081 | -0.07 | -0.01 | 0.60 | -0.15 | | -0.05 | 0.77 | 0.71 | 0.18 |
| 2082 | -0.28 | -0.56 | 0.94 | -0.33 | -0.04 | 0.07 | -0.42 | -0.09 | -0.15 |
| 2083 | 0.04 | -0.09 | -0.08 | 0.07 | -0.32 | 0.29 | -0.67 | 0.09 | -0.12 |
| 2084 | -0.14 | -0.27 | -0.49 | 0.27 | 0.11 | 0.04 | 1.55 | -0.28 | |
| 2087 | -0.18 | -0.23 | 0.25 | 0.76 | 0.05 | | -1.56 | -0.07 | -0.62 |
| 2101 | -0.45 | -0.51 | 0.01 | -0.25 | -0.11 | | 1.01 | 0.01 | 0.47 |
| 2102 | 0.07 | -0.48 | 0.05 | | 0.73 | -0.25 | 0.54 | 0.43 | -0.58 |
| 2111 | -0.24 | -0.28 | -0.71 | -0.13 | -0.05 | -0.24 | -0.45 | -0.88 | -0.32 |
| 2112 | 0.31 | 0.38 | 0.19 | 0.58 | 0.24 | | -0.63 | | -0.29 |

**Table S3**. **Analysis results of systematic bias between visits in dorsolateral prefrontal cortex.**

Variations between the first and second visits were calculated. A negative variation means that there is a decrease in the second visit and is highlighted in red. A positive variation means that there is an increase in the second visit and is highlighted in green. A variation of “0.00” means that there is no change between visits and is highlighted in gray. There was not a systematic increase or decrease observed between visits. This suggests that there is not a technique batch effect that may affect the longitudinal analysis results. An empty cell means the data was excluded from analysis due to low quality. Abbreviations: GABA indicates γ-aminobutyric acid; Gln, glutamine; Glu, glutamate; GSH, glutathione; Lac, lactate; NAA, N-acetylaspartate; NAAG, N-acetylaspartyl glutamate; mI, myo-inositol; tCho, phosphocholine plus glycerophosphocholine.

| **Subject** | **GSH** | **GABA** | **Glu** | **Gln** | **tCho** | **Lac** | **mI** | **NAA** | **NAAG** |
|---|---|---|---|---|---|---|---|---|---|
| 1001 | 0.22 | -0.34 | -1.30 | 1.05 | -0.09 | -0.08 | 0.03 | -0.13 | -0.04 |
| 1005 | 0.65 | -0.66 | 2.37 | 0.09 | 0.15 | | 0.55 | 1.32 | 0.33 |
| 1006 | -0.44 | 0.03 | -0.65 | -0.07 | 0.03 | -1.64 | -0.33 | -0.35 | |
| 1007 | -0.04 | 0.14 | -0.45 | 0.20 | -0.31 | | -0.84 | -0.40 | -0.04 |
| 1011 | 0.43 | 0.32 | 1.74 | 0.41 | 0.54 | 0.16 | 0.97 | 0.45 | 0.03 |
| 1014 | -0.28 | -0.08 | -0.61 | -0.58 | -0.09 | 0.51 | -0.46 | -0.19 | -0.10 |
| 1018 | -0.20 | -0.11 | 0.08 | 0.06 | -0.43 | | -0.35 | 0.68 | 0.07 |
| 1022 | 0.44 | 0.68 | 1.67 | 0.38 | 0.31 | 0.38 | 1.32 | 1.78 | 0.09 |
| 1026 | 0.12 | -0.27 | 0.34 | -0.28 | 0.01 | -0.05 | 0.81 | -0.66 | 0.51 |
| 1032 | | 0.23 | 0.03 | 0.61 | 0.00 | 0.13 | -0.23 | 0.39 | -0.09 |
| 1033 | | | 0.16 | -0.36 | 0.29 | | 0.08 | 0.18 | |
| 1038 | 0.26 | 0.28 | -0.19 | 0.50 | 0.06 | 0.33 | -1.26 | -0.93 | -0.20 |
| 1039 | -0.15 | -0.30 | -0.64 | -0.70 | -0.10 | 0.83 | -0.44 | -0.53 | 0.22 |
| 1042 | 0.30 | 0.07 | -0.20 | 0.68 | 0.09 | 0.01 | 0.13 | -0.16 | -0.39 |
| 1044 | -0.05 | -0.24 | -0.56 | -1.15 | -0.20 | -1.32 | -1.49 | -0.62 | 0.14 |
| 1047 | 0.18 | 1.14 | 1.62 | 0.27 | -0.02 | | -0.84 | 0.86 | 0.10 |
| 1050 | -0.20 | -0.23 | -0.70 | 0.16 | -0.24 | -0.16 | -0.53 | -0.08 | -0.43 |
| 1051 | 0.12 | 0.06 | -0.55 | -0.17 | 0.15 | 0.19 | 0.37 | 0.13 | 0.27 |
| 1052 | 0.04 | -0.44 | 0.32 | 0.37 | -0.03 | | -0.60 | 1.32 | |
| 1055 | 0.15 | -0.12 | -0.92 | 0.21 | 0.20 | -0.35 | 0.62 | -0.33 | -0.77 |
| 1056 | 0.05 | -0.38 | -0.73 | -0.43 | -0.07 | 0.14 | -0.09 | 0.34 | -0.12 |
| 1070 | 1.05 | 0.73 | 1.68 | 0.14 | 0.71 | -0.29 | 1.95 | 2.45 | 0.21 |
| 1082 | -0.07 | 0.16 | 0.59 | 0.40 | | | 0.04 | -1.33 | |
| 1085 | | 0.04 | 0.40 | 0.32 | 0.10 | | -0.55 | -0.06 | 0.15 |
| 1090 | -0.22 | | -0.33 | 0.08 | | -0.18 | -0.25 | -1.22 | 0.49 |
| 1109 | -0.13 | 0.16 | -0.83 | 0.07 | -0.08 | -1.14 | -0.28 | | 0.48 |
| 1118 | 0.26 | 0.01 | 0.38 | 0.41 | 0.10 | | -0.08 | -0.48 | 0.00 |
| 1119 | 0.33 | 0.06 | 0.18 | -0.30 | -0.16 | 0.07 | 0.25 | 0.41 | 0.01 |
| 1122 | 0.30 | -0.82 | -0.47 | 0.16 | 0.10 | | -0.85 | 0.44 | |

(continued on next page)

| | | | | | | | | | |
|---|---|---|---|---|---|---|---|---|---|
| 1132 | 0.36 | 0.28 | 1.28 | 0.43 | 0.38 | | 1.51 | -0.17 | 0.33 |
| 1136 | 0.24 | -0.13 | -0.24 | -0.08 | 0.04 | | 0.04 | -0.84 | |
| 2001 | 0.01 | -0.15 | -0.75 | -0.17 | -0.07 | 0.52 | 0.01 | 0.11 | -0.57 |
| 2002 | -0.18 | 0.76 | 1.38 | 0.47 | 0.23 | 0.10 | 0.22 | 0.72 | 0.51 |
| 2003 | -0.39 | 0.13 | -1.29 | 0.03 | -0.25 | -0.80 | -0.73 | -1.30 | |
| 2005 | 0.11 | 0.39 | 0.61 | 0.21 | -0.07 | -0.36 | 0.00 | -0.23 | 0.04 |
| 2007 | 0.12 | 0.11 | 0.35 | 0.34 | 0.35 | 0.18 | 0.25 | -0.10 | 0.64 |
| 2009 | 0.22 | 0.39 | 0.22 | | 0.50 | -0.35 | 0.27 | 0.03 | 0.15 |
| 2010 | -0.10 | -0.14 | -1.10 | -0.43 | 0.11 | | -0.44 | -0.20 | 0.63 |
| 2014 | 0.19 | 0.06 | 0.51 | 0.11 | 0.21 | 0.12 | 0.14 | 0.45 | -0.16 |
| 2016 | 0.30 | 0.13 | 0.03 | 0.59 | 0.19 | -0.18 | 0.21 | -0.04 | -0.27 |
| 2019 | -0.21 | 0.11 | 0.02 | 0.38 | 0.07 | -0.45 | 0.15 | -0.31 | 0.39 |
| 2022 | -0.07 | 0.43 | 1.42 | 0.61 | 0.19 | -1.34 | -0.42 | -0.90 | -0.37 |
| 2023 | -0.14 | 0.21 | -0.29 | -0.05 | -0.12 | -0.23 | -0.46 | -0.14 | -0.09 |
| 2024 | 0.26 | 0.20 | 0.50 | 0.28 | -0.01 | 0.03 | -0.04 | -0.26 | 0.06 |
| 2026 | 0.23 | 0.33 | 0.47 | 0.09 | 0.11 | 1.01 | 0.49 | -1.14 | |
| 2028 | -0.03 | -0.10 | 0.57 | 0.44 | 0.13 | -0.33 | 0.29 | 0.71 | |
| 2029 | | -0.93 | 0.37 | 0.85 | 0.18 | | 0.98 | -0.85 | 0.23 |
| 2032 | -0.01 | -0.38 | -0.48 | -0.32 | -0.18 | 0.09 | 0.18 | 0.55 | -0.53 |
| 2038 | | -0.25 | 0.08 | 0.28 | 0.17 | -0.28 | 0.57 | 0.06 | 0.41 |
| 2041 | 0.13 | -0.26 | -1.23 | -0.69 | -0.08 | | | -1.56 | |
| 2042 | 0.24 | -0.35 | 0.52 | -0.01 | 0.09 | | -1.14 | -1.44 | |
| 2047 | 0.07 | -0.02 | -0.11 | -0.06 | -0.23 | -0.04 | -0.57 | -0.38 | -0.27 |
| 2048 | -0.11 | 0.16 | 0.50 | 0.40 | 0.10 | -0.48 | -0.61 | 0.44 | 0.18 |
| 2050 | -0.09 | | -0.37 | -0.08 | 0.08 | -0.85 | -0.88 | -0.73 | |
| 2052 | 0.01 | -0.10 | -0.38 | | -0.25 | | -0.38 | 0.33 | 0.32 |
| 2053 | 0.00 | 0.03 | 0.56 | 0.35 | -0.10 | -0.39 | -0.42 | 0.39 | |
| 2056 | 0.08 | -0.13 | -0.41 | -0.22 | 0.00 | 0.12 | -0.20 | -0.88 | -0.14 |
| 2057 | 0.59 | -0.23 | 3.67 | | 0.35 | | 1.12 | | |
| 2062 | | -0.60 | 0.93 | 0.49 | 0.13 | | 0.48 | 0.19 | |
| 2066 | -0.19 | -0.17 | -0.01 | -0.28 | | -0.70 | 0.08 | | -0.41 |
| 2067 | -0.23 | -0.34 | -0.38 | 0.02 | | | | 1.22 | |
| 2068 | -0.01 | -0.53 | -1.02 | -0.31 | 0.00 | | -0.30 | -0.95 | -0.09 |
| 2069 | -0.18 | -0.08 | -0.97 | -0.03 | -0.07 | | -0.30 | -0.19 | |
| 2070 | 0.41 | -0.01 | 0.07 | 0.30 | 0.26 | | 0.64 | 0.46 | -0.46 |
| 2071 | | 0.62 | 0.09 | | 0.32 | -0.88 | 0.29 | -0.05 | |
| 2073 | | | -0.49 | -0.04 | | -0.16 | -0.02 | -0.60 | 0.29 |
| 2078 | 0.16 | | -0.29 | -0.61 | -0.15 | 0.24 | -0.05 | -0.35 | 0.17 |
| 2080 | -0.09 | -0.57 | -0.48 | 0.16 | 0.03 | -0.10 | -0.34 | -0.13 | |
| 2081 | -0.33 | 0.23 | -0.92 | -0.18 | -0.32 | | -0.58 | -0.63 | |
| 2082 | 0.16 | -0.77 | -0.42 | -0.41 | 0.20 | | -0.16 | 0.80 | -0.23 |
| 2083 | 0.19 | 0.29 | 0.06 | 0.39 | -0.16 | | -0.91 | -0.41 | |
| 2087 | 0.06 | 0.05 | 0.14 | -0.14 | -0.16 | | 0.11 | 0.22 | 0.16 |
| 2099 | | -0.30 | -0.26 | -0.63 | | -0.26 | -0.35 | -0.51 | -0.27 |
| 2101 | 0.45 | 0.50 | 0.30 | -0.01 | | | -0.16 | 0.07 | |
| 2102 | 0.23 | -0.04 | -0.06 | -0.18 | 0.27 | -0.12 | 0.18 | -0.63 | 0.21 |
| 2111 | | -0.58 | -1.05 | -0.86 | -0.20 | | -1.18 | -2.47 | 1.06 |
| 2112 | 0.50 | -0.05 | 0.44 | -0.46 | -0.05 | | -0.10 | 0.10 | -0.57 |

**Table S4**. **Analysis results of systematic bias between visits in centrum semiovale.**

Variations between the first and second visits were calculated. A negative variation means that there is a decrease in the second visit and is highlighted in red. A positive variation means that there is an increase in the second visit and is highlighted in green. A variation of "0.00" means that there is no change between visits and is highlighted in gray. There was not a systematic increase or decrease observed between visits. This suggests that there is not a technique batch effect that may affect the longitudinal analysis results. An empty cell means the data was excluded from analysis due to low quality. Abbreviations: GABA indicates γ-aminobutyric acid; Gln, glutamine; Glu, glutamate; GSH, glutathione; Lac, lactate; NAA, N-acetylaspartate; NAAG, N-acetylaspartyl glutamate; mI, myo-inositol; tCho, phosphocholine plus glycerophosphocholine.

| **Subject** | **GSH** | **GABA** | **Glu** | **Gln** | **tCho** | **Lac** | **mI** | **NAA** | **NAAG** |
|---|---|---|---|---|---|---|---|---|---|
| 1001 | 0.19 | 0.06 | -0.48 | 0.84 | -0.27 | -0.09 | 0.04 | -0.73 | -0.24 |
| 1005 | -0.19 | 0.03 | -0.25 | -0.10 | 0.09 | 0.40 | -0.35 | -0.47 | 0.76 |
| 1006 | -0.13 | -0.07 | -0.92 | -0.30 | -0.09 | -0.42 | 0.03 | 0.54 | 0.85 |
| 1007 | 0.04 | -0.17 | -0.58 | -0.11 | -0.07 | 0.24 | 0.46 | -0.27 | 0.15 |
| 1011 | 0.06 | 0.18 | 0.13 | 0.14 | 0.32 | 0.10 | 0.64 | 0.29 | 0.30 |
| 1012 | | -0.16 | -0.03 | -0.27 | -0.24 | -0.04 | -1.48 | -0.82 | -0.08 |
| 1014 | -0.35 | -0.01 | 0.26 | 0.39 | -0.43 | -0.09 | -0.21 | -0.85 | 0.08 |
| 1022 | 0.04 | -0.13 | -0.41 | 0.10 | 0.09 | 0.16 | 0.67 | 0.26 | 0.12 |
| 1026 | 0.50 | 0.53 | 1.21 | | 0.28 | 0.44 | 0.59 | -0.23 | 0.36 |
| 1032 | 0.34 | 0.16 | -0.08 | 0.53 | -0.13 | 0.05 | -0.14 | 0.33 | -0.23 |
| 1033 | 0.05 | -0.20 | 0.06 | -0.45 | -0.32 | -0.02 | -0.25 | -0.24 | -0.44 |
| 1038 | 0.19 | -0.02 | 0.93 | | 0.03 | 0.35 | -0.98 | 0.50 | 0.21 |
| 1039 | -0.03 | -0.68 | -0.52 | -0.55 | -0.25 | | -0.13 | 0.07 | -0.16 |
| 1042 | -0.03 | 0.10 | 0.53 | 0.54 | | 0.23 | -0.20 | -0.61 | 0.13 |
| 1044 | 0.07 | -0.19 | -0.30 | | 0.23 | | 0.03 | 0.75 | 0.12 |
| 1047 | | 0.49 | | 1.13 | | 0.39 | | | |
| 1050 | 0.19 | 0.27 | 0.05 | 0.16 | 0.06 | -0.07 | 0.38 | 0.58 | 0.06 |
| 1051 | 0.14 | 0.05 | -0.51 | 0.09 | 0.06 | 0.15 | 0.40 | -0.03 | 0.04 |
| 1052 | 0.09 | -0.05 | 0.46 | -0.05 | -0.06 | -0.26 | 0.30 | -0.06 | -0.19 |
| 1055 | -0.23 | -0.39 | -0.04 | 0.12 | 0.24 | 0.07 | 0.77 | -0.85 | 0.12 |
| 1056 | -0.12 | | -0.57 | -0.13 | 0.08 | -0.06 | 0.70 | 0.29 | 0.15 |
| 1070 | 0.53 | 0.85 | 0.75 | | 0.35 | 0.37 | 1.51 | 0.67 | 0.83 |
| 1082 | -0.18 | -0.24 | -0.62 | -0.39 | 0.07 | 0.02 | -0.57 | -0.27 | -0.26 |
| 1085 | -0.24 | -0.38 | -1.16 | -0.12 | -0.12 | -0.21 | 0.04 | -0.46 | 0.03 |
| 1090 | | -0.11 | -1.09 | -0.23 | -0.02 | 0.04 | -0.84 | -1.53 | 0.29 |
| 1091 | -0.26 | 0.32 | -0.45 | -0.66 | 0.27 | 0.61 | 0.10 | -0.64 | |
| 1105 | 0.35 | 0.08 | 0.08 | 0.13 | -0.15 | 0.06 | -0.43 | -1.55 | -0.35 |
| 1109 | 0.30 | 0.05 | -0.39 | -0.54 | 0.30 | 0.56 | 0.36 | 0.56 | 0.23 |
| 1118 | -0.16 | 0.02 | -0.23 | -0.41 | -0.01 | -0.05 | 0.02 | 0.08 | -0.06 |

(continued on next page)

| | | | | | | | | | |
|---|---|---|---|---|---|---|---|---|---|
| 1122 | 0.07 | 0.28 | | 0.43 | -0.16 | | -1.09 | 0.12 | 0.19 |
| 1132 | 0.50 | | | 0.32 | 0.54 | -0.20 | 1.69 | | 0.67 |
| 1136 | | 0.08 | -0.08 | 0.18 | 0.00 | -0.06 | -0.18 | -0.29 | 0.11 |
| 2001 | 0.04 | 0.18 | 0.27 | 0.13 | -0.08 | -0.18 | -0.30 | 0.00 | 0.05 |
| 2002 | -0.13 | 0.02 | -0.44 | 0.35 | -0.25 | -0.17 | -0.79 | -0.71 | -0.08 |
| 2003 | -0.12 | 0.10 | -0.07 | -0.26 | -0.03 | -0.17 | 0.38 | -0.43 | -0.08 |
| 2005 | 0.04 | 0.06 | 0.52 | -0.17 | -0.22 | 0.50 | -0.54 | 0.02 | -0.44 |
| 2007 | -0.05 | -0.03 | 0.55 | 0.27 | 0.02 | -0.18 | 0.10 | -0.13 | -0.15 |
| 2009 | -0.16 | -0.23 | -0.24 | -0.47 | -0.06 | -0.06 | 0.11 | -0.38 | -0.31 |
| 2010 | -0.05 | 0.02 | 0.02 | -0.22 | 0.05 | -0.21 | -0.60 | -0.51 | 0.12 |
| 2014 | 0.09 | 0.30 | 0.69 | 0.06 | -0.01 | 0.02 | -0.24 | -0.08 | -0.08 |
| 2016 | 0.13 | 0.03 | 0.67 | 0.07 | -0.13 | 0.00 | 0.05 | -0.39 | -0.21 |
| 2019 | 0.26 | -0.13 | -0.16 | 0.34 | 0.10 | 0.32 | 0.48 | 0.11 | 0.15 |
| 2022 | -0.31 | -0.16 | -0.58 | -0.20 | 0.05 | -0.04 | -0.22 | -0.85 | |
| 2023 | -0.08 | -0.08 | -0.95 | -0.11 | 0.08 | 0.04 | -0.22 | 0.29 | 0.26 |
| 2024 | 0.16 | 0.14 | 0.15 | 0.23 | 0.10 | 0.13 | -0.02 | -0.02 | -0.17 |
| 2026 | | 0.06 | -0.16 | 0.10 | | -0.06 | 0.00 | -0.25 | -0.46 |
| 2028 | 0.19 | 0.34 | 0.80 | 0.02 | 0.15 | 0.19 | 0.43 | 0.47 | 0.29 |
| 2029 | 0.00 | 0.01 | 0.60 | 0.11 | -0.23 | 0.12 | -0.31 | -0.70 | -0.48 |
| 2032 | -0.02 | -0.08 | 0.20 | 0.17 | 0.01 | -0.15 | 0.32 | -0.44 | -0.35 |
| 2038 | 0.21 | 0.35 | -0.53 | -0.17 | 0.09 | 0.04 | 0.73 | 0.86 | 0.31 |
| 2041 | 0.01 | 0.01 | 0.18 | -0.37 | -0.20 | -0.19 | 0.13 | 0.06 | -0.38 |
| 2042 | 0.00 | | 0.14 | 0.02 | 0.08 | 1.12 | 0.52 | 0.25 | -0.13 |
| 2047 | 0.07 | 0.14 | 0.82 | | -0.08 | 0.24 | 0.12 | 0.33 | -0.34 |
| 2048 | -0.03 | -0.08 | -0.01 | | 0.25 | 0.04 | -0.32 | -0.10 | 0.29 |
| 2050 | 0.11 | 0.04 | 0.73 | 0.26 | 0.15 | -0.33 | 0.13 | 0.88 | -0.08 |
| 2052 | 0.13 | -0.22 | -0.73 | 0.19 | -0.05 | 0.34 | 0.12 | -0.32 | -0.06 |
| 2053 | -0.01 | -0.09 | -0.45 | -0.06 | -0.10 | -0.02 | -0.71 | -0.70 | -0.11 |
| 2056 | -0.23 | -0.05 | -0.63 | -0.17 | 0.03 | -0.02 | -0.53 | -0.54 | 0.04 |
| 2057 | 0.19 | -0.41 | 0.15 | -0.20 | -0.42 | 0.04 | -0.35 | -0.73 | -1.01 |
| 2062 | -0.32 | 0.11 | -0.11 | 0.00 | 0.11 | | -0.24 | -0.54 | 0.67 |
| 2066 | | -0.18 | 0.11 | 0.20 | 0.01 | -0.43 | 0.03 | -0.10 | -0.07 |
| 2067 | -0.21 | 0.11 | -0.29 | -0.02 | 0.12 | 0.01 | 0.04 | -0.38 | 0.26 |
| 2068 | -0.04 | 0.01 | -0.39 | -0.24 | 0.03 | 0.08 | 0.06 | 0.08 | 0.02 |
| 2069 | -0.20 | -0.01 | 0.17 | 0.13 | -0.25 | 0.07 | 0.13 | 0.11 | -0.19 |
| 2070 | 0.03 | 0.27 | 0.32 | 0.03 | 0.07 | 0.03 | 0.37 | 0.55 | 0.16 |
| 2071 | 0.02 | | -0.64 | -0.09 | 0.01 | 0.24 | 0.38 | 0.21 | |
| 2073 | 0.35 | -0.07 | -0.19 | 0.14 | -0.07 | -0.11 | 0.32 | 0.30 | -0.23 |
| 2075 | 0.00 | -0.11 | -0.06 | | -0.03 | | 0.17 | 0.20 | -0.37 |
| 2078 | -0.20 | | -0.10 | -0.08 | 0.04 | 0.86 | 0.04 | -0.08 | 0.35 |
| 2080 | -0.14 | 0.18 | 0.27 | 0.32 | -0.07 | 0.26 | -0.47 | -0.18 | 0.12 |
| 2081 | 0.19 | -0.03 | -0.20 | -0.20 | -0.14 | 0.22 | -0.42 | -0.13 | -0.18 |
| 2082 | 0.14 | 0.28 | -0.43 | | -0.03 | 0.54 | -0.03 | 0.82 | -0.25 |
| 2083 | 0.01 | 0.29 | 0.29 | 0.51 | 0.05 | 0.37 | 0.25 | -0.17 | 0.34 |
| 2084 | 0.17 | -0.15 | -0.93 | -0.40 | 0.10 | 0.54 | 0.13 | -0.03 | 0.11 |
| 2087 | 0.22 | | 0.27 | 0.01 | -0.24 | | -0.07 | -0.33 | -0.80 |
| 2099 | | -0.06 | -0.37 | 0.03 | 0.16 | -0.05 | 0.69 | | 0.20 |
| 2102 | -0.31 | | -0.25 | -0.08 | -0.05 | 0.33 | -0.46 | -0.41 | 0.00 |
| 2111 | 0.00 | 0.03 | 0.08 | -0.22 | -0.08 | -0.16 | | -0.35 | -0.15 |
| 2112 | 0.01 | 0.04 | 0.24 | 0.05 | | 0.01 | -0.13 | | |

**Table S5**. **Analysis results of systematic bias between visits in orbitofrontal region.**

Variations between the first and second visits were calculated. A negative variation means that there is a decrease in the second visit and is highlighted in red. A positive variation means that there is an increase in the second visit and is highlighted in green. A variation of "0.00" means that there is no change between visits and is highlighted in gray. There was not a systematic increase or decrease observed between visits. This suggests that there is not a technique batch effect that may affect the longitudinal analysis results. An empty cell means the data was excluded from analysis due to low quality. Abbreviations: GABA indicates γ-aminobutyric acid; Gln, glutamine; Glu, glutamate; GSH, glutathione; Lac, lactate; NAA, N-acetylaspartate; NAAG, N-acetylaspartyl glutamate; mI, myo-inositol; tCho, phosphocholine plus glycerophosphocholine.

| **Subject** | **GSH** | **GABA** | **Glu** | **Gln** | **tCho** | **Lac** | **mI** | **NAA** | **NAAG** |
|---|---|---|---|---|---|---|---|---|---|
| 1006 | 0.32 | -0.30 | -1.04 | -0.26 | -0.29 | 0.04 | 0.10 | -0.32 | -0.16 |
| 1007 | 0.16 | -0.14 | -0.41 | 0.23 | -0.12 | 0.08 | 0.54 | 0.13 | -0.16 |
| 1011 | -0.02 | 0.22 | 0.37 | 0.22 | -0.11 | 0.27 | -0.53 | -0.21 | 0.34 |
| 1012 | 0.14 | -0.21 | 0.27 | 0.33 | 0.06 | -0.03 | -0.11 | -0.21 | |
| 1014 | -0.20 | -1.76 | -1.73 | -0.39 | -0.69 | 0.05 | -1.39 | -1.37 | -0.77 |
| 1018 | 0.23 | 0.08 | 0.41 | 0.06 | -0.06 | | 0.49 | 0.17 | 0.25 |
| 1022 | 0.13 | -0.58 | -0.71 | 0.65 | -0.21 | | -0.78 | -0.42 | 1.03 |
| 1026 | -0.02 | 0.45 | 0.70 | 0.45 | -0.17 | -0.18 | 0.20 | -0.34 | -0.07 |
| 1032 | 0.22 | -0.02 | -0.28 | 0.33 | 0.05 | 0.33 | -0.01 | 0.17 | -0.22 |
| 1033 | -0.26 | -0.39 | -0.89 | -0.67 | -0.41 | | 0.15 | -0.16 | -0.57 |
| 1038 | 0.13 | 0.16 | -0.18 | -0.31 | 0.32 | | -2.32 | 0.46 | |
| 1039 | -0.60 | -0.22 | 0.04 | 0.57 | -0.69 | 0.09 | -0.40 | -0.42 | |
| 1040 | 0.01 | 0.69 | 0.45 | 0.52 | 0.21 | 0.26 | -1.13 | -0.28 | 0.24 |
| 1042 | 0.45 | 0.29 | 0.41 | 0.31 | 0.06 | -0.23 | 1.51 | -0.67 | |
| 1051 | 0.40 | -0.76 | -0.35 | -0.06 | 0.35 | 0.31 | 0.71 | 0.56 | |
| 1052 | 0.02 | -0.08 | 0.38 | -0.23 | 0.16 | -0.28 | -0.14 | 0.35 | 0.42 |
| 1056 | -0.23 | 0.10 | 0.80 | 0.06 | -0.12 | | -1.16 | 0.67 | -0.20 |
| 1070 | 0.58 | 0.64 | 0.08 | 0.14 | 0.06 | 0.05 | 0.58 | 0.23 | 1.35 |
| 1082 | -0.10 | 0.32 | -0.06 | -0.38 | -0.10 | | -0.10 | -0.56 | -0.06 |
| 1085 | -0.16 | 0.55 | 0.84 | 0.60 | 0.11 | | 1.47 | 0.08 | |
| 1109 | | -0.82 | -1.52 | | -0.17 | | 0.19 | -1.07 | |
| 1119 | 0.26 | | 0.26 | 0.62 | 0.04 | 0.08 | -0.50 | 0.29 | 0.54 |
| 1122 | -0.31 | -0.41 | -1.55 | 0.54 | -0.22 | | -0.26 | -0.95 | |
| 1132 | 0.13 | -0.21 | 0.38 | 0.19 | -0.10 | -0.19 | -0.08 | -0.30 | |
| 1136 | -0.07 | -0.01 | -0.21 | 0.45 | 0.05 | -0.59 | -0.25 | 0.10 | |
| 2001 | 0.45 | -0.78 | -2.03 | -0.94 | 0.53 | -0.13 | -0.45 | 0.02 | 1.21 |
| 2002 | -0.05 | 0.10 | -0.58 | 0.20 | 0.00 | 0.13 | -1.07 | -0.37 | 0.51 |
| 2005 | -0.20 | -0.40 | -0.49 | 0.25 | -0.02 | -0.13 | 0.40 | -0.03 | 0.20 |
| 2007 | 0.28 | 0.61 | -0.02 | 0.16 | 0.22 | | -0.08 | 0.36 | -0.53 |

(continued on next page)

| | | | | | | | | | |
|---|---|---|---|---|---|---|---|---|---|
| 2009 | -0.41 | -0.64 | -0.86 | -0.49 | 0.34 | | -1.31 | -0.47 | 1.14 |
| 2010 | -0.11 | -0.72 | -0.28 | -0.17 | -0.17 | | 0.25 | -0.55 | 0.05 |
| 2014 | -0.76 | -0.94 | 1.52 | 0.00 | 0.01 | | 3.37 | -0.93 | |
| 2016 | -0.11 | 1.30 | 0.94 | 0.30 | 0.29 | | -2.12 | -0.38 | 0.83 |
| 2019 | -0.06 | -0.10 | -0.34 | 0.14 | -0.01 | 0.03 | 0.61 | -0.65 | 0.10 |
| 2022 | 0.20 | 0.67 | 0.54 | 0.16 | -0.16 | 0.13 | -0.12 | -0.12 | -0.10 |
| 2023 | 0.39 | 0.18 | 0.30 | 0.09 | -0.18 | 0.19 | 0.11 | 0.58 | -0.44 |
| 2024 | 0.03 | -0.82 | -1.21 | -1.03 | -0.09 | -0.34 | 0.37 | -0.52 | |
| 2026 | 0.32 | -0.05 | 0.61 | 0.07 | 0.26 | -0.32 | 0.36 | 0.95 | 0.19 |
| 2029 | | 0.69 | 0.33 | -0.40 | 0.10 | -0.15 | 0.35 | 1.19 | |
| 2032 | 0.27 | -0.35 | 0.65 | 0.30 | -0.19 | -0.19 | 1.47 | -0.54 | 0.01 |
| 2038 | 0.14 | -0.44 | -0.57 | 0.12 | -0.38 | | -0.49 | -0.07 | 0.05 |
| 2042 | 0.11 | -0.37 | -0.81 | 0.14 | | | 0.13 | -0.70 | -0.21 |
| 2047 | | 0.22 | 0.09 | | -0.02 | -0.10 | -0.71 | | |
| 2048 | -0.13 | -0.23 | -0.13 | -0.57 | -0.15 | -0.44 | -0.93 | -0.46 | -0.70 |
| 2050 | 0.49 | 0.13 | 0.60 | 0.02 | 0.07 | -0.06 | 0.72 | 0.64 | -0.38 |
| 2052 | 0.16 | -0.67 | -0.04 | 0.28 | 0.38 | | 0.06 | 0.63 | -0.06 |
| 2053 | 0.10 | -0.39 | -1.13 | -0.03 | -0.01 | -0.48 | -0.49 | -0.11 | -0.32 |
| 2056 | 0.32 | 0.14 | -0.32 | -0.15 | 0.13 | 0.64 | -0.16 | | -0.20 |
| 2057 | 0.38 | -0.09 | 0.09 | 0.51 | 0.10 | | 0.74 | 1.40 | -0.32 |
| 2062 | -0.05 | 0.18 | 0.22 | -0.25 | 0.16 | -0.49 | -0.21 | 0.09 | 0.21 |
| 2066 | 0.15 | 0.21 | 0.63 | 0.33 | | -0.23 | -0.05 | -0.38 | -0.64 |
| 2067 | 0.02 | -0.66 | -0.66 | -0.39 | 0.10 | | -0.32 | -0.12 | |
| 2068 | -0.09 | 0.10 | 0.59 | -0.19 | -0.03 | 0.14 | 1.08 | -0.51 | 0.57 |
| 2069 | -0.03 | -0.23 | -0.53 | 0.26 | -0.28 | -0.61 | 0.27 | -0.74 | |
| 2070 | -0.03 | -0.62 | | -0.35 | -0.44 | -0.37 | 0.51 | -0.34 | -0.61 |
| 2071 | -0.41 | -0.38 | -0.24 | | 0.11 | -0.28 | -0.07 | -0.47 | 0.03 |
| 2073 | 0.39 | -0.34 | -0.67 | -0.26 | 0.04 | | 0.96 | 0.04 | -0.30 |
| 2075 | 0.43 | 0.18 | 1.10 | | 0.16 | -0.79 | 1.32 | 0.55 | |
| 2078 | 0.05 | 0.45 | 0.92 | 0.30 | -0.29 | | -0.37 | 0.61 | 0.48 |
| 2082 | 0.24 | 0.35 | 0.78 | 0.15 | -0.12 | 0.54 | -0.20 | 0.01 | -0.27 |
| 2083 | 0.16 | -0.11 | 0.93 | 0.35 | -0.30 | -0.21 | -0.17 | 0.44 | 0.13 |
| 2084 | | 0.42 | 0.08 | 0.33 | 0.04 | -0.50 | 0.02 | -0.36 | |
| 2087 | 0.08 | -0.36 | | 0.26 | -0.11 | 0.36 | 0.86 | 0.74 | -0.52 |
| 2099 | -0.34 | -0.40 | -0.18 | -0.26 | -0.01 | | 0.56 | 0.16 | -0.21 |
| 2102 | -0.16 | 0.02 | -0.90 | -0.61 | -0.04 | -0.21 | -0.29 | -0.59 | 0.17 |
| 2111 | 0.41 | 0.48 | 1.11 | | -0.01 | | 0.76 | 0.06 | 0.25 |
| 2112 | -0.51 | -0.63 | -0.38 | -1.05 | | -0.15 | -1.01 | -0.97 | -0.30 |

**Table S6. Analysis results for return bias in healthy controls**.

We compared the metabolite levels in 5 brain regions between the full cohort and the longitudinal cohort. There was not a significant difference observed in any metabolite. This suggests that there is not a bias in healthy controls that returned for the longitudinal study. Abbreviations: GABA indicates γ-aminobutyric acid; Gln, glutamine; Glu, glutamate; GSH, glutathione; Lac, lactate; NAA, N-acetylaspartate; NAAG, N-acetylaspartyl glutamate; mI, myo-inositol; tCho, phosphocholine plus glycerophosphocholine; ACC, anterior cingulate cortex; Thal, thalamus; DLPFC, dorsolateral prefrontal cortex; CSO, centrum semiovale; and OFR, orbital frontal cortex.

| **region** | **metabolite** | **p-value** |
|---|---|---|
| ACC | GSH | 0.52 |
| ACC | GABA | 0.78 |
| ACC | Glu | 0.84 |
| ACC | Gln | 0.93 |
| ACC | tCho | 0.26 |
| ACC | Lac | 0.69 |
| ACC | mI | 0.47 |
| ACC | NAA | 0.79 |
| ACC | NAAG | 0.91 |
| DLPFC | GSH | 0.62 |
| DLPFC | GABA | 0.81 |
| DLPFC | Glu | 0.84 |
| DLPFC | Gln | 0.80 |
| DLPFC | tCho | 0.37 |
| DLPFC | Lac | 0.98 |
| DLPFC | mI | 0.86 |
| DLPFC | NAA | 0.79 |
| DLPFC | NAAG | 0.39 |
| Thal | GSH | 0.70 |
| Thal | GABA | 0.92 |
| Thal | Glu | 0.07 |
| Thal | Gln | 0.64 |
| Thal | tCho | 0.38 |
| Thal | Lac | 0.97 |
| Thal | mI | 0.56 |
| Thal | NAA | 0.67 |
| Thal | NAAG | 0.84 |

(continued on next page)

| | | |
|---|---|---|
| CSO | GSH | 0.48 |
| CSO | GABA | 0.85 |
| CSO | Glu | 0.58 |
| CSO | Gln | 0.75 |
| CSO | tCho | 0.70 |
| CSO | Lac | 0.13 |
| CSO | mI | 0.64 |
| CSO | NAA | 0.90 |
| CSO | NAAG | 0.92 |
| OFR | GSH | 0.16 |
| OFR | GABA | 0.30 |
| OFR | Glu | 0.76 |
| OFR | Gln | 0.49 |
| OFR | tCho | 0.75 |
| OFR | Lac | 0.76 |
| OFR | mI | 0.91 |
| OFR | NAA | 0.62 |
| OFR | NAAG | 0.89 |

**Table S7. Analysis results for return bias in first episode psychosis patients**.

We compared the metabolite levels in 5 brain regions between the full cohort and the longitudinal cohort. There was not a significant difference observed in any metabolite. This suggests that there is not a bias in patients that returned for the longitudinal study. Abbreviations: GABA indicates γ-aminobutyric acid; Gln, glutamine; Glu, glutamate; GSH, glutathione; Lac, lactate; NAA, N-acetylaspartate; NAAG, N-acetylaspartyl glutamate; mI, myo-inositol; tCho, phosphocholine plus glycerophosphocholine; ACC, anterior cingulate cortex; Thal, thalamus; DLPFC, dorsolateral prefrontal cortex; CSO, centrum semiovale; and OFR, orbital frontal cortex.

| region | metabolite | p-value |
|---|---|---|
| ACC | GSH | 0.19 |
| ACC | GABA | 0.66 |
| ACC | Glu | 0.32 |
| ACC | Gln | 0.53 |
| ACC | tCho | 0.67 |
| ACC | Lac | 0.35 |
| ACC | mI | 0.57 |
| ACC | NAA | 0.52 |
| ACC | NAAG | 0.74 |
| DLPFC | GSH | 0.67 |
| DLPFC | GABA | 0.95 |
| DLPFC | Glu | 0.71 |
| DLPFC | Gln | 0.86 |
| DLPFC | tCho | 0.89 |
| DLPFC | Lac | 0.88 |
| DLPFC | mI | 0.92 |
| DLPFC | NAA | 0.75 |
| DLPFC | NAAG | 0.62 |
| Thal | GSH | 0.36 |
| Thal | GABA | 0.55 |
| Thal | Glu | 0.62 |
| Thal | Gln | 0.58 |
| Thal | tCho | 0.84 |
| Thal | Lac | 0.36 |
| Thal | mI | 0.37 |
| Thal | NAA | 0.71 |
| Thal | NAAG | 0.66 |

(continued on next page)

| | | |
|---|---|---|
| CSO | GSH | 0.91 |
| CSO | GABA | 0.60 |
| CSO | Glu | 0.50 |
| CSO | Gln | 0.53 |
| CSO | tCho | 0.84 |
| CSO | Lac | 0.70 |
| CSO | mI | 0.74 |
| CSO | NAA | 0.39 |
| CSO | NAAG | 0.87 |
| OFR | GSH | 0.79 |
| OFR | GABA | 0.66 |
| OFR | Glu | 0.52 |
| OFR | Gln | 0.35 |
| OFR | tCho | 0.88 |
| OFR | Lac | 0.74 |
| OFR | mI | 0.79 |
| OFR | NAA | 0.39 |
| OFR | NAAG | 0.70 |

**Figure S1**. **Impact of confounding factors on anterior cingulate cortex (ACC) metabolite levels in the first episode psychosis (FEP) patient group.**

In the analysis comparing FEP patients and healthy controls (HC), we adjusted for the confounding effects of age, gender, race, and smoking status. The patient group may be influenced by intrinsic factors (such as duration of illness (DOI)) and extrinsic factors (such as chlorpromazine (CPZ) equivalent dose), which cannot be adjusted in two-group comparisons since they don't apply to HC. To evaluate the impact of these factors, including demographic variates, we performed linear regression with gender and race (which don't change over time) as covariates. Absolute (abs) T values were used to quantitively measure the impact. The red dotted line shows the cutoff of significance (the T value corresponding to a p-value of 0.05). Abbreviations: GABA indicates γ-aminobutyric acid; Glu, glutamate; GSH, glutathione; NAA, N-acetylaspartate; mI, myo-inositol; and tCho, phosphocholine plus glycerophosphocholine.

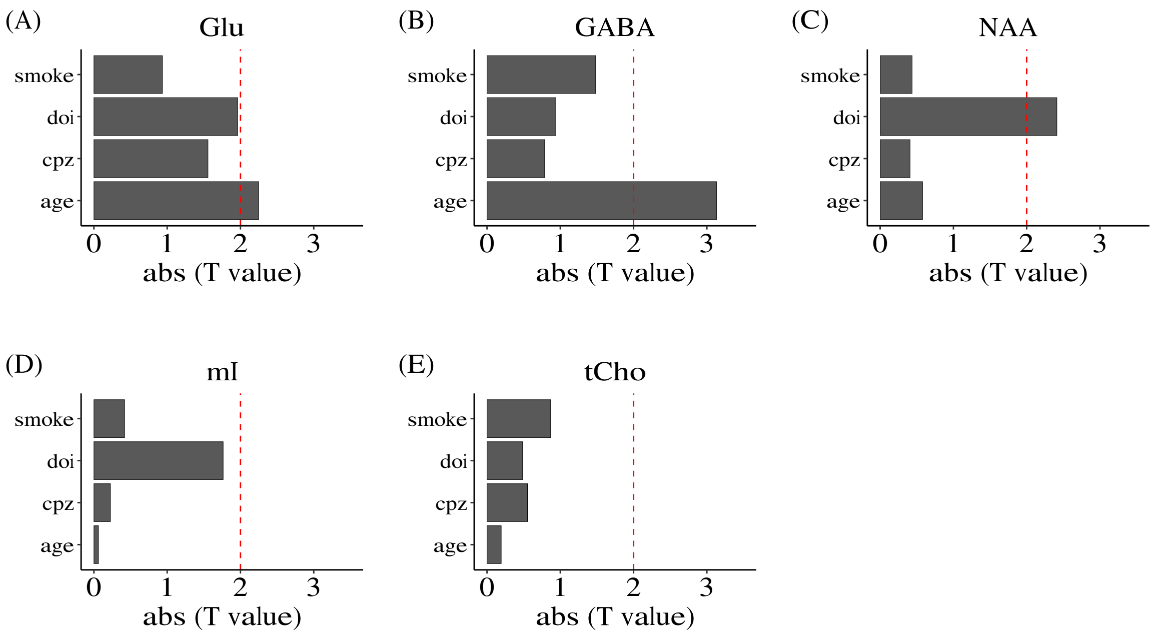
